# Nature portrayed in images in Dutch Brazil: Tracing the sources of the plant woodcuts in the *Historia Naturalis Brasiliae* (1648)

**DOI:** 10.1101/2022.10.04.510872

**Authors:** Mireia Alcantara-Rodriguez, Tinde Van Andel, Mariana Françozo

## Abstract

By the mid-seventeenth century, images of natural elements that originated in Dutch Brazil circulated in Europe. These were often included in art collections (the *Libri Picturati*), and natural history treatises (the *Historia Naturalis Brasiliae* and the *India Utriesque re Naturale et Medica*). The plant woodcut images in these books constituted (icono) type specimens and played a significant role in the transmission of (botanical) knowledge. We present a systematic analysis of their origins by cross-referencing the visual and textual sources related to Dutch Brazil. To do so, we used the scientific identifications of the portrayed plants and digital archival material. The plant woodcuts accounted for 529 images, which correspond to 426 taxa. We created a PDF booklet to visualize the correlations of the woodcuts with the *Libri Picturati* and other visual sources. Substantial differences in the visual-making methodology exist between the two treatises (1648, 1658). Overall, availability, economy, and the Indigenous Tupi-based plant names that accompanied the images were crucial when arranging the sources, as well as portraying as much botanical information as possible.

Freshly picked, living plants, and dried branches, fruits, and seeds were used to represent the megadiverse Brazilian flora, even when these belonged to species originating from other regions. Despite not being recognized for their contribution, Indigenous Brazilians and enslaved Africans were essential in the visual knowledge-making processes that later resulted in these natural history collections. As several sources remain lost and many histories yet untold, further archival studies and collaborative projects are pertinent to reveal the missing pieces of this conundrum.

## 1 Introduction

### 1.1 Nature portrayed in the early modern period: Dutch Brazil

By the mid-seventeenth century, images of natural elements that originated in Dutch Brazil circulated in Europe, often included in art collections and natural history treatises. The term ‘Dutch Brazil’ corresponds to the northeastern part of Brazil that was colonized by the Dutch West Indian Company (WIC) between 1630 and 1654 after they overtook it from the Portuguese. Appointed by the WIC, count Johan Maurits of Nassau-Siegen was the governor-general in Dutch Brazil from 1636 to 1644. He commissioned naturalist George Marcgrave, physician Willem Piso, and painters Albert Eckhout and Frans Post, among others, to portray and document the environment encountered in the colony. The outcomes of the team assembled by Johan Maurits included, among others, the treatises *Historia Naturalis Brasiliae* (HNB) (1) and *India Utriesque re Naturale et Medica* (IURNM) (2), botanical specimens in Marcgrave’s herbarium, and a set of plant illustrations, drawings, and sketches divided into three collections: *Theatrum Rerum Naturalium Brasiliae* (*Theatrum*), *Miscellanea Cleyeri* (*Misc. Cleyeri*), and the *Libri Principis* (also known as *Handbooks* or *Manuais*). The botanical imagery of these sources is the object of this study.

### 1.2 Background to the visual and textual repertoire of Dutch Brazil

The origin, arrangement, and destination of all these materials are varied. The HNB was commissioned by Johan Maurits after his return from Brazil, when he handed over the field notes of Piso and Marcgrave and several drawings on plants and animals to Johannes de Laet, WIC director, cartographer, and ultimately editor of the HNB. De Laet organized these drawings and ordered a few woodcut images to be made based on Marcgrave’s herbarium, as becomes clear by his commentaries (3). The HNB was published in 1648 by Elzevier in Amsterdam and Hackium in Leiden, becoming an authoritative work on Brazilian and tropical flora and fauna for the upcoming centuries (4,5). It contains references to classical naturalists, such as Dioscorides and Theophrastus, and to Renaissance and contemporary scholars (e.g., Francisco Hernández, Nicolas Monardes, and Carolus Clusius), whose manuscripts and early images of American flora influenced the work of Marcgrave and Piso (5,6).

Based on the field notes, De Laet produced a preliminary draft of the HNB (7), which included 15 plant drawings, 11 proof-woodcuts, and 366 plant descriptions – often with the word *Icon* next to them (4). This manuscript could be De Laet’s first attempt to arrange Marcgrave’s species. The word *Icon* could correspond to field drawings or watercolor illustrations and oil paintings from the expedition (8), or living plants collected in Brazil (4). The manuscript was later purchased by the slaveholder, physician, and collector Hans Sloane (1660-1753) and it is now part of the Sloane manuscript collection at the British Library. Most of Marcgrave’s specimens ended up in Copenhagen in 1653 and are now kept at the University herbarium (C).

In 1644, Johan Maurits, Piso, Ekckout, Post, and other members of the count’s crew returned to the Low Countries, but Marcgrave never did. In 1643, he was sent to Angola, where he died shortly upon his arrival (9). De Laet died in 1649, a year after the publication of the HNB. Piso, discontent with the arrangement of the HNB, edited its content and published a modified version in 1658 (2).

The Brazilian imagery had another destiny. In 1652, Johan Maurits sent several oil-based illustrations and drawings of flora, fauna, and Indigenous and African peoples from Dutch Brazil as a diplomatic gift to Frederick William, Elector of Brandenburg from 1640 to 1688. He passed them to his court physician Christian Mentzel, who organized the oil paintings in a bound collection (the *Theatrum*). Mentzel included Johan Maurits’s words in the preface of the *Theatrum*, emphasizing that ‘these images aimed to reproduce Brazilian nature as perfectly as possible’ (10: 18). The elector was also gifted with two bound volumes of watercolors (the *Libri Principis*), and a few sketches and oil paintings, which were bound in 1757 (the *Misc. Cleyeri*) (4). In the nineteenth century, these Brazilian collections were incorporated into a larger collection known as the *Libri Picturati*, which was housed in the Preussische Staatsbibliothek (the present-day *Staatsbibliothek* in Berlin, Germany), until this library was evacuated during World War II (11). The *Libri Picturati* were considered lost until zoologist Peter Whitehead located them at the Jagiellonian Library in Krakow, Poland (4,12).

### 1.3 Behind the images: previous and present research

Due to its undeniable historical value, several studies on the Brazilian images in the *Libri Picturati* have been conducted, focusing on its content, composition, and authorship (4,20,21,11,13–19). Scholars have argued that the *Libri Picturati* images served as models for the woodcuts of the HNB (4,24,25,11,12,14–16,18,22,23). Correlations between the *Theatrum* illustrations and the woodcuts had been established for the birds (26) and the animals in general – also including the *Libri Principis* (15,25). A few woodcuts seem to be based on Post’s drawings (27). Brazilian botanists Pickel (6) and Almeida (5) suggested the borrowing of images from published treatises, such as those by Clusius or De Laet. Some woodcuts were made in the Dutch Republic after the specimens collected by Marcgrave (8). Additionally, several woodcuts were based on Marcgrave’s drawings, as De Laet indicated in the preface of the HNB (1), as well as Piso in his book (2). In 1640, Marcgrave wrote to De Laet from Brazil that he had made 350 pencil drawings of plants and several more of animals (28). The few pencil sketches found in De Laet’s manuscript could be part of those (4,8,22), but most of Marcgrave’s original drawings do not exist anymore or have not been located yet. Beyond doubt, the study of the woodcuts embedded in these natural history books is of great relevance, as they play a significant role in the transmission of knowledge (29) and they constituted (icono) type specimens: they accompanied the first descriptions of individual species against which all later individuals were compared (4). As the intriguing question of which plant images were used as the basis for the woodcuts remained unanswered, we present a systematic analysis of their origins. We focused on the botanical images included in the HNB, the IURNM, the *Libri Picturati* Brazilian collection, De Laet’s manuscript, and Marcgrave’s herbarium specimens to 1) analyze the correlations between the woodcuts and the other visual sources from Dutch Brazil, and 2) trace back the remaining sources that were used to create the woodcuts. By applying our botanical image analysis to these historical collections, we provide an overview of how the visual material was used in the composition of seventeenth century natural history treatises on Brazil. In doing so, we add insights into the processes of visual knowledge-making and botanical practices in the early modern period.

## 2 Materials and Methods

To analyze the correlations among the historical visual sources, we built a database in FileMaker Pro with all woodcuts and illustrations organized by species and created a spreadsheet document with the background information. Our database contained all woodcut images present in Marcgrave’s and Piso’s books (1,2) their correspondent images for the same species in Marcgrave’s herbarium (collected between 1638 and 1643), the *Libri Picturati*, and other visual sources. We used a digital-colored copy of the HNB located in Leiden University (the Netherlands) ([HNB Leiden University library](https://digitalcollections.universiteitleiden.nl/view/item/1535938)) (Fig 1) and the digital images of Marcgrave’s herbarium in C ([Marcgrave’s herbarium](https://samlinger.snm.ku.dk/en/dry-and-wet-collections/botany/general-herbarium/the-marcgrave-herbarium/)). For the IURNM, we used the copy kept at the Missouri Botanical Garden ([IURNM copy Missouri](https://archive.org/details/mobot31753002909064)). The *Libri Picturati* illustrations were provided as digital images by the Jagiellonian library, of which the *Miscellanea Cleyeri* ([Misc. Cleyeri](https://jbc.bj.uj.edu.pl/dlibra/doccontent?id=197455)) and the *Libri Principis* ([Libri Principis](https://jbc.bj.uj.edu.pl/dlibra/publication/193892/edition/183824/content)) are publicly available.

**Fig 1.**
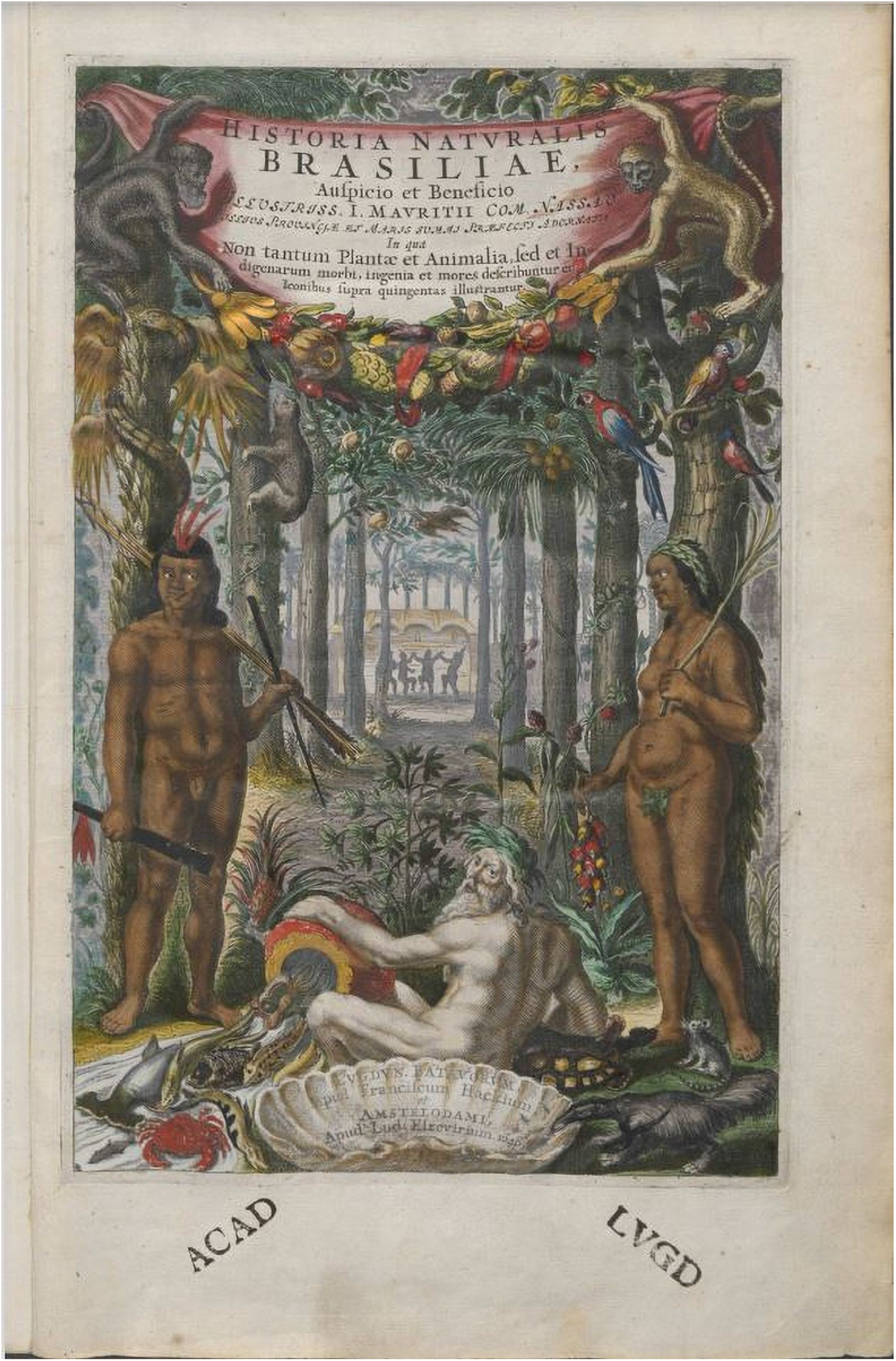
Frontispiece of the *Historia Naturalis Brasiliae* (Marcgrave and Piso, 1648). Colored copy kept at the Library of Leiden University (the Netherlands). Available at https://digitalcollections.universiteitleiden.nl/view/item/1535938#page/25/mode/1up.

To study the connections between this material and the woodcut images, we compared the woodcuts with the images included in De Laet’s manuscript in the Sloane collection (7). We associated the plant descriptions that included the term ‘Icon’ and the drawings glued to the manuscript to their corresponding species in the HNB. The digitized scans of De Laet’s manuscript were provided by the Manuscript Department of the British Library, London.

To compare woodcuts, vouchers, and plant illustrations, we cross-referenced all textual and visual sources for each plant species by using our recent botanical identifications for the HNB, the IURNM, and the *Libri Picturati* (30,31). Then, we assembled the images from multiple sources that belonged to the same species to visualize their (dis-)similarities by using FileMaker Pro. Due to the larger number of woodcuts and the many duplicates in Piso’s work (2,32), we started our analysis with Marcgrave’s (1648) chapters on plants (3). We systematized the information with database entries on the page numbers of the woodcuts in the HNB and IURNM, vernacular name(s), botanical identifications, and the origin of these woodcuts. We first identified the woodcuts that correlated at the species level with Marcgrave’s herbarium and the *Libri Picturati*. For the latter, we analyzed the degree of similarity among the images. We distinguished four categories: 1) very similar (woodcut and illustration share [almost] the same features), 2) moderately similar (they bear a great resemblance but do not share the same features), 3) slightly similar (woodcuts share some characteristics but not as many as in the previous category), and 4) different (not enough similar features between images to assume any correlation).

For the woodcut images that did not resemble the visual sources mentioned above, we checked the works of Renaissance and early modern scholars that included engravings similar to the plant images in the HNB and IURNM. To navigate this vast corpus of literature, we first checked the HNB and IURNM for references to scholars who worked with tropical flora, such as Hernández (33), Monardes (34), or Clusius (35–37). We narrowed this search by consulting Almeida (5), who systematized those citations per plant species. For Clusius, we searched for corresponding images in Ubrizsy and Heniger (38), who listed the American plants portrayed by this botanist. For the woodcuts whose sources were still unknown, we searched for their species name on Plantillustrations.org (plantillustrations.org). This open-access site offers a large digital collection of HD botanical illustrations through time. We retrieved some of these flora illustrations for our database.

Finally, we checked phenological and other botanical characters in modern photos (including herbarium vouchers) and added them to the database to show how these plants appear in nature. All plant images were retrieved from Creative Commons (creativecommons.org), Flickr (flickr.com), Plants of the World Online (powo.science.kew.org), Flora do Brasil 2020 (floradobrasil.jbrj.gov.br), Species Link (specieslink.net), and the Global Biodiversity Information Facility (gbif.org).

## 3 Results

### 3.1 Origin of the *Historia Naturalis Brasiliae* (1648) woodcuts

There is a total of 301 plant woodcuts in the HNB (1). The background information of all woodcut images (species, family, sources, page numbers, author, etc.) is listed in Supporting information S1. A complete overview of all the visual sources arranged by their correlated species, taking as the main reference the woodcuts in the HNB is provided in Supplementary information S2. Fig 2 shows how we linked a woodcut in the HNB to several other visual sources.

**Fig 2.**
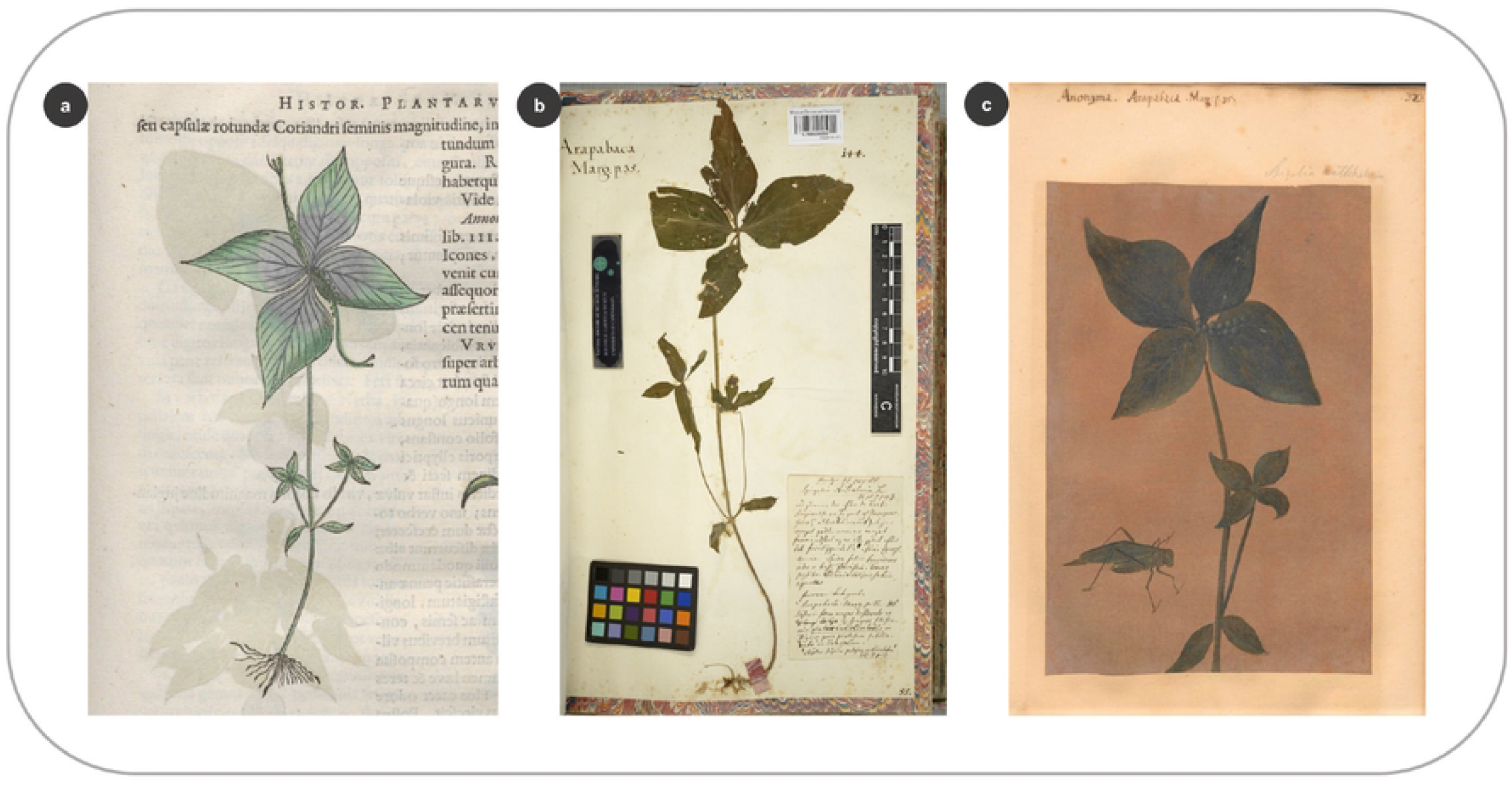
Linking a woodcut in the HNB to other visual sources from Dutch Brazil. (a) Woodcut of *Spighelia anthelmia* L. in the HNB (Marcgrave 1648: 35) (b) Specimen of the same species in Marcgrave’s herbarium (f. 55) (c) *S. anthelmia* depicted as an oil painting in the *Theatrum* (f. 323).

Several woodcuts in Marcgrave’s chapters on plants are repeated in Piso’s chapter (39) because De Laet re-used them for the species that were mentioned by both authors (Fig 3). In the HNB, a total of nine species are represented by two different woodcuts, while two are depicted by three different ones. For six woodcuts we were unable to identify the species depicted (Fig 3).

**Fig 3.**
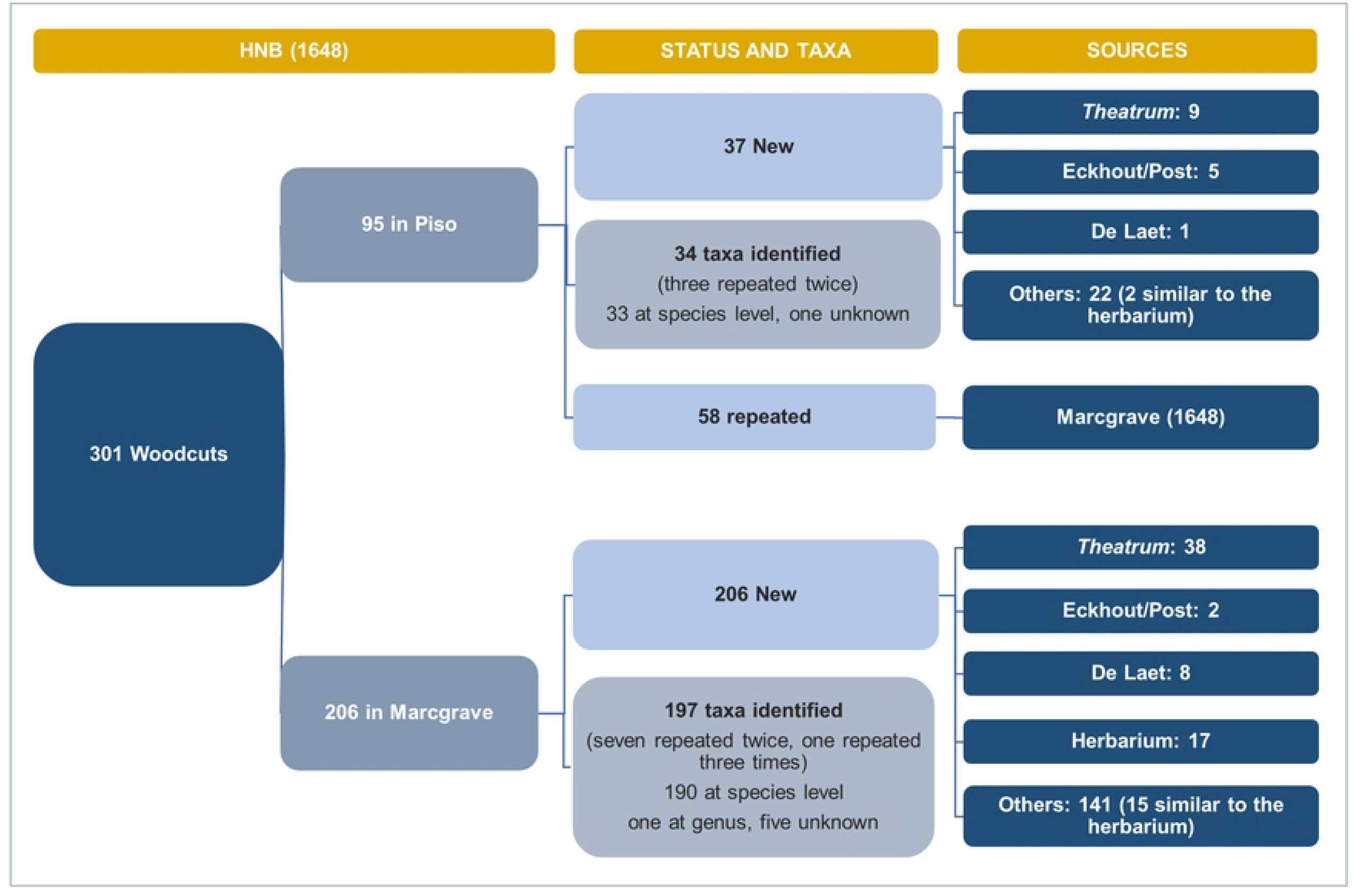
Tracing the sources of the plant woodcuts in the HNB. Flowchart showing the organization of the woodcut images in the HNB, the number of plant taxa, and the sources used to create the woodcuts.

We traced approximately one-third (84 woodcuts) of the 243 unique woodcuts in the HNB (Marcgrave’s woodcuts plus Piso’s unique ones) to the *Libri Picturati*, Marcgrave’s herbarium, or other known sources (Fig 3). Over two-thirds (159 woodcuts) of the plant images did not correspond to any of these visual sources.

#### 3.1.1 Theatrum Rerum Naturalium

From the entire Brazilian collection of the *Libri Picturari*, the HNB plant woodcuts mostly match with the *Theatrum* illustrations. One exception is the woodcut that represents *Canna indica* L., which is very similar to the one in *Misc. Cleyeri* (in reversed format). Most of the plant woodcuts in the HNB, however, represent species not included in the *Theatrum* (Fig 3), even though over half of the species in common with the *Theatrum* (78 species in Marcgrave and 15 spp. in Piso), bear resemblance in different degrees to the plant illustrations, while the other half are different (Figs 4 and 5). A small proportion (15%, 7 spp.) of the overlapping images between the HNB and the *Libri Picturati* (47 spp.) are reversed (example in Fig 5b).

**Fig 4.**
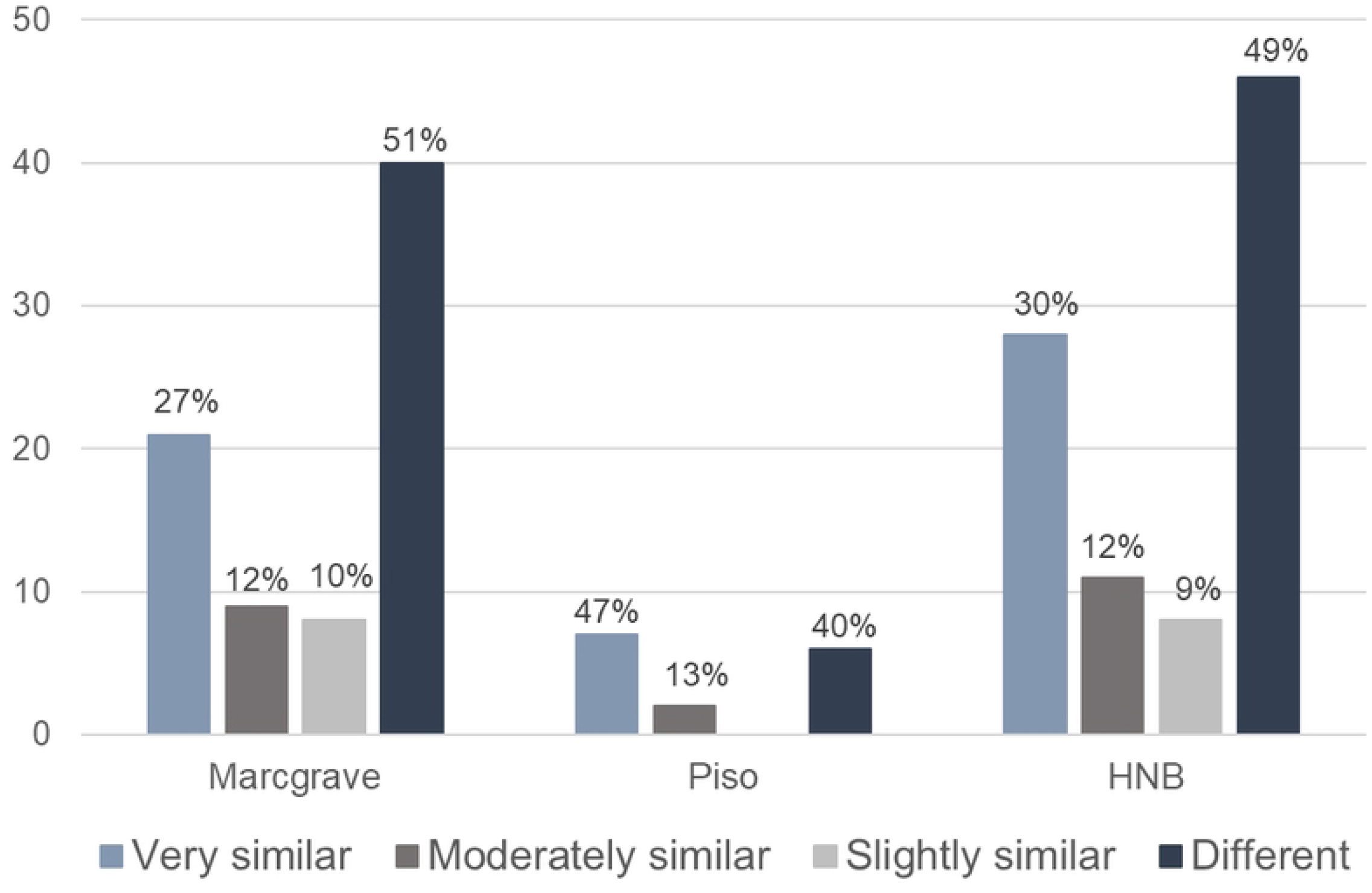
Comparing the plant images portrayed in the HNB and the *Theatrum* for the same species. Similarities between the botanical illustrations in the *Theatrum* and the woodcuts depicted in Marcgrave’s chapters on plants, Piso’s chapter on medicinal plants, and the whole repertoire of botanical woodcuts in the HNB (Marcgrave and Piso 1648).

**Fig 5.**
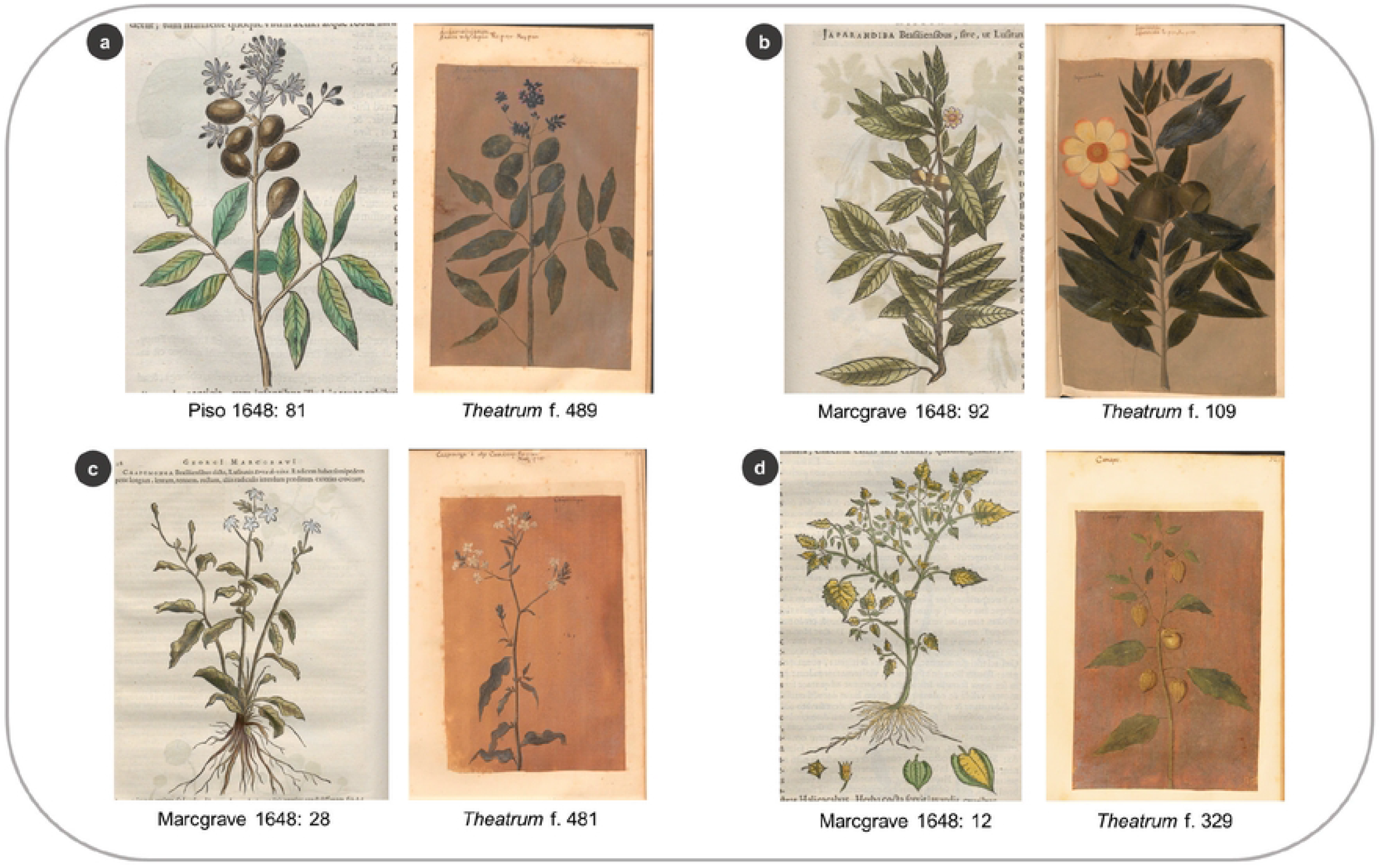
Examples of similarity degrees between the woodcuts in the HNB and the *Theatrum* illustrations. (a) Very similar: *Andira fraxinifolia* Benth. (b) Moderately similar: *Gustavia augusta* L. (c) Slightly similar: *Plumbago zeylanica* L. (d) Different: *Physalis pubescens* L.

#### 3.1.2 Marcgrave’s herbarium

Out of the 143 species preserved in Marcgrave’s herbarium, 92 (64%) are described in Marcgrave and Piso’s books, but only 74 are represented by a woodcut image in the HNB. According to Andrade-Lima et al. (8), 17 woodcuts for the HNB were produced using Marcgrave’s herbarium vouchers (Fig 3, S1). De Laet himself explicitly mentioned in the HNB that 15 woodcuts were made after these specimens (3). In addition, *Galphimia brasiliensis* (L.) A.Juss. and *Spondias mombin* L. were also made after Marcgrave’s exsiccates (S2: 303, 449), although De Laet did not indicate this in the HNB (8). By comparing the taxa shared between the HNB and the herbarium, we found that the woodcuts of 25 species bear no resemblance to the vouchers; hence these were probably made after other sources. For 17 species in the herbarium, we cannot infer whether the specimens were used to design the woodcuts, because they are poorly preserved or consist of a few plant parts (a single leaf or sterile branches). Nevertheless, 17 specimens bear similarities to the woodcut images (Fig 3), so they could have been used as models to elaborate the drawings that were later used to carve the woodcuts (Fig 6).

**Fig 6.**
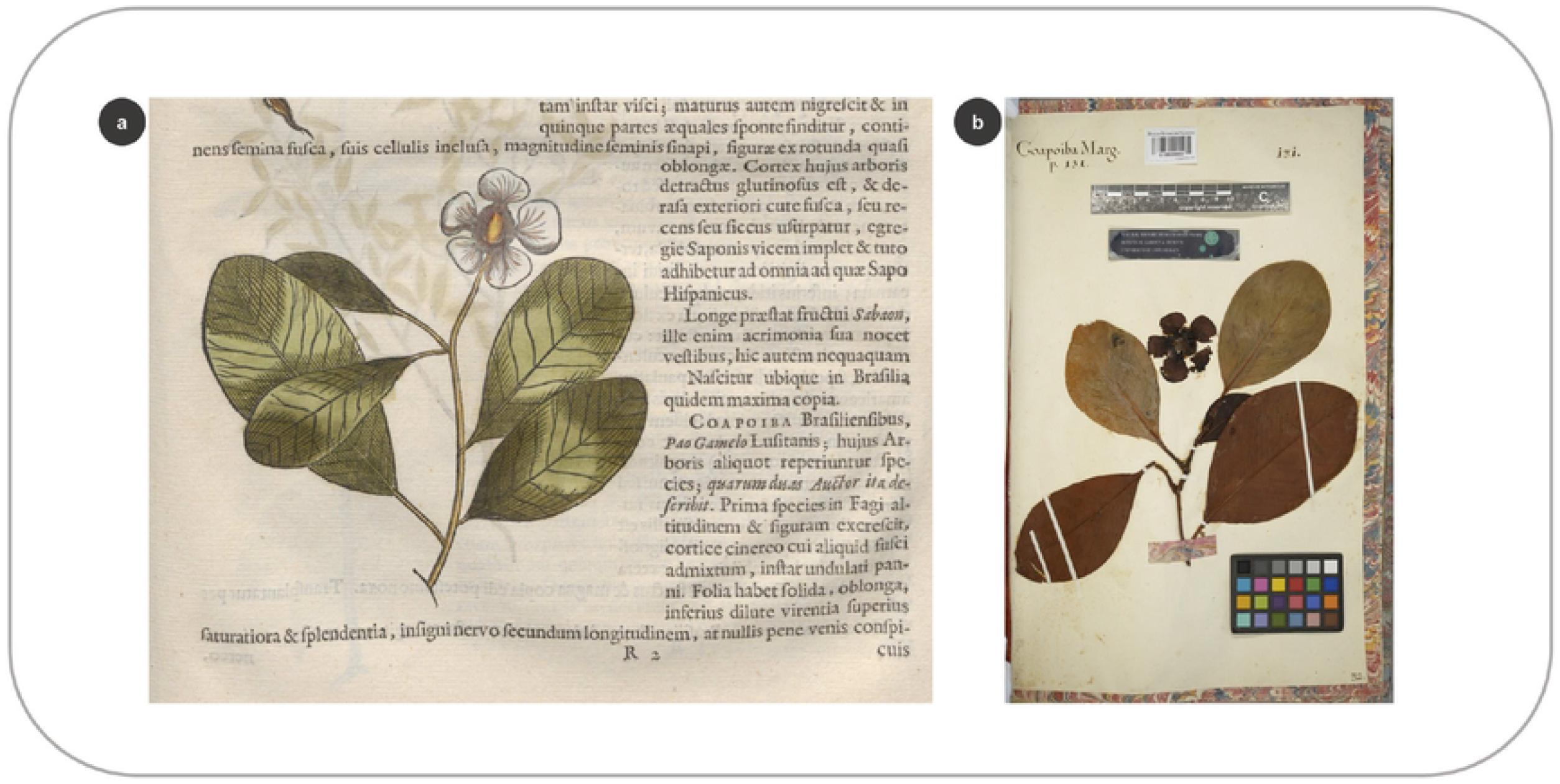
Correlation between a woodcut in the HNB and the specimen collected by Marcgrave. (a) Woodcut of *Clusia nemorosa* G.Mey. in the HNB (Marcgrave 1648: 131) (b) Specimen of *C. nemorosa* in Marcgrave’s herbarium (f. 32).

#### 3.1.3 Other traceable visual sources

De Laet tried to find ways to provide images of the plants described in the HNB when these were lacking – as occurred with the vouchers collected by Marcgrave. De Laet received a pod of *Mimosa pigra* L. from his friends in Brazil, which he commissioned to have designed at its natural size, even though “the painter hardly represented its elegance” (3: 74). While editing the HNB, De Laet also obtained seeds from Brazil of a plant called *Micambe Angolensibus*.

He planted them and observed how the plant grew and flowered in the summer of 1646, but also saw it die in October (3). This was *Cleome gynandra* L., an African weed introduced to Brazil via the trans-Atlantic slave trade and eaten by enslaved Africans as a leafy vegetable (3: 10). De Laet tried to obtain a drawing from the living plant in the Dutch Republic, but he did not manage to do so, as the tropical plant perished during the winter frost (3).

We found that eight woodcuts were based on images previously published by De Laet in his books on the colonized territories of the Americas (40,41). For instance, De Laet mentioned that the woodcut of *Bromelia karatas* L. depicted on p. 111 in Piso’s chapter on medicinal plants (39) was drawn at its natural scale after he received the fruit from Brazil depicted on page 614 (40). In turn, De Laet (re)used three of the woodcuts from his books, which he copied after images of related flora published by Clusius (36,37) (S2). De Laet often cited Clusius, Monardes, and other Renaissance authors to compare the Brazilian plants to known European or American species.

The artists that joined Johan Maurits’s crew in Brazil also played a role in the making of the images. Three woodcuts included in Piso (39) depict how sugarcane (*Saccharum officinarum* L.) and cassava (*Manihot esculenta* Crantz) were processed in the mills and ovens of the colony by enslaved African labor. These are thought to have been designed by Frans Post (4,27). We found similar scenes in Barlaeus’ 1647 publication about Johan Maurits’ endeavors in the colony (42), which were indeed drawn by Post and appeared embedded in the maps designed by Marcgrave in Brazil (S2: 1-5). The woodcut of *Cereus jamacaru* DC. also bears a strong resemblance to the cactus portrayed by Post in ‘View of the Rio São Francisco Brazil with Fort Maurits and Capibara’ (S2: 437).

Post’s contemporary Albert Eckhout included at least three plants in his still-life paintings and portraits that are similar to the woodcuts in the HNB (S1, S2). One example is *Ipomoea pes-caprae* (L.) R. Br, of which the woodcut in Marcgrave’s chapter in the HNB does not match any of the known sources (Fig 7a), but the same species in Piso (Fig 7b) strongly resembles the creeping plant in Eckhout’s portrait known as ‘African man’ (Figs 7c, 7d).

**Fig 7.**
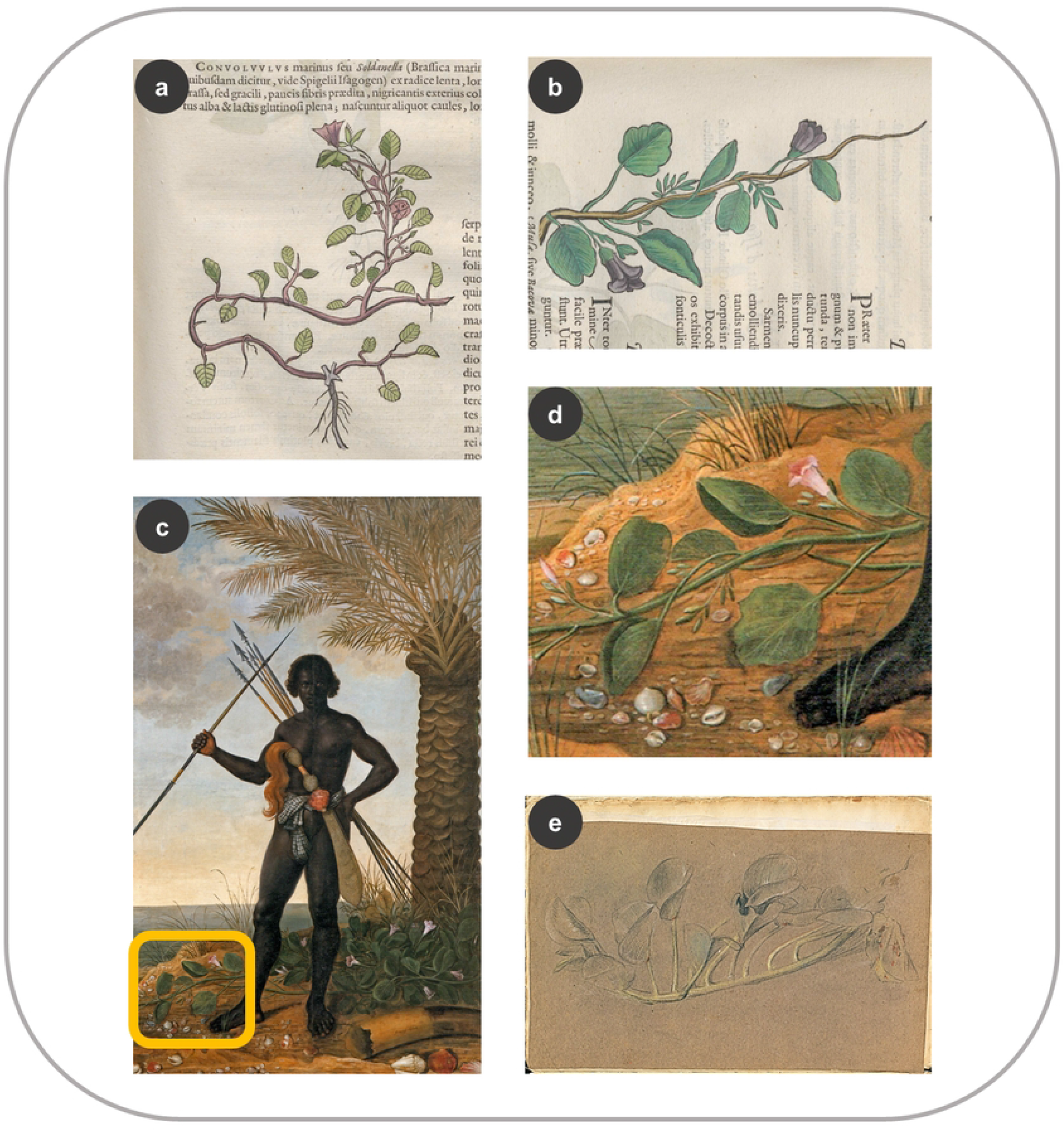
Similarities between the HNB and Eckhout’s paintings. (a) Woodcut of *Ipomoea pes-caprae* (Marcgrave 1648: 51) (b) *I. pes-caprae* in Piso (1648: 103) (c) the same species on the left bottom corner in Eckhout’s painting (National Museum of Denmark) (d) close-up of *I. pes-caprae* in Eckhout’s painting (e) Sketch of the same species in the *Misc. Cleyeri* (f. 12v).

There is a crayon sketch of the same species in the *Misc. Cleyeri* (Fig 7e), but it neither resembles the woodcut (Fig 7b) nor the vine in the painting (Fig 7d) – as is often the case with the sketches in the *Misc. Cleyeri* and Eckhout’s paintings (16,21).

#### 3.1.4 Remaining sources of the HNB woodcuts

Most of the HNB woodcut images (67%, 163 woodcuts) could not be traced to either the *Libri Picturati*, Marcgrave’s herbarium, Eckhout and Post’s paintings, or De Laet. The question is then whether these woodcuts were made by Marcgrave himself. Interestingly, considerable mistakes are present in several of these woodcuts. On page 137, the banana bunch (*Musa* × *paradisiaca* L.) emerges from the middle part of the trunk, instead of from the center (3). The *Anda* tree (*Joannesia princeps* Vell.) on page 110 bears large, bell-shaped flowers (3) compared to the smaller and compound inflorescences characteristic of the Euphorbiaceae family (S2: 377-378). It is unlikely that a trained botanist like Marcgrave would have approved these morphological errors in his botanical drawings (6).

In Marcgrave’s chapter on trees, De Laet indicated for *Jetaiba* (*Hymenaea* cf. *courbaril* L.) that ‘more details about this tree were to be found in Piso, to whom we owe this figure’ (3: 101). De Laet referred to the woodcut in Piso, which is the same as in Marcgrave’s chapter (39: 60). It is unclear whether Piso made some of the plant drawings since he was not trained in botany and art like his naturalist colleague. We know that some woodcuts were borrowed from Marcgrave’s sketches during their expeditions (2: 107, 43: 249), while others were made *ad vivum* (after life) by a painter traveling with him in the *sertão*, the dry hinterland of northeastern Brazil (preface in (2: 2, 43: 8).

### 3.2 Origin of the woodcuts in the *India Utriusque re Naturale et Medica* (1658)

The background information of all woodcut images (species, family, sources, page numbers, author, etc.) is listed in Supporting information S3, and the associated images are displayed in Supporting information S4. Piso reused many woodcuts from the HNB, which are shown in S2. Hence, to avoid repetitions, we only added in S4 those plant woodcuts that are different. In most cases, the woodcuts from Marcgrave’s chapters on plant species that were not described as used by humans are lacking in the IURNM.

### 3.1.1 New and reused woodcut images

In the IURNM, there are in total 228 woodcuts, eight of which are depicted twice (Fig 8). Hence, 220 different woodblocks must have been used to complete the botanical part of Piso’s *solo* work. These are distributed in Piso’s chapter IV, which is the equivalent to Marcgrave’s chapters on flora and the chapter on medicinal plants by Piso (44), and in Piso’s chapter V, which is the equivalent to the chapter on venoms and antidotes in the HNB (45). Within the 59 identified taxa (Fig 8), we found a mushroom, a sponge, and a coral, which do not belong to the plant kingdom but were classified as such by seventeenth century scholars.

**Fig 8.**
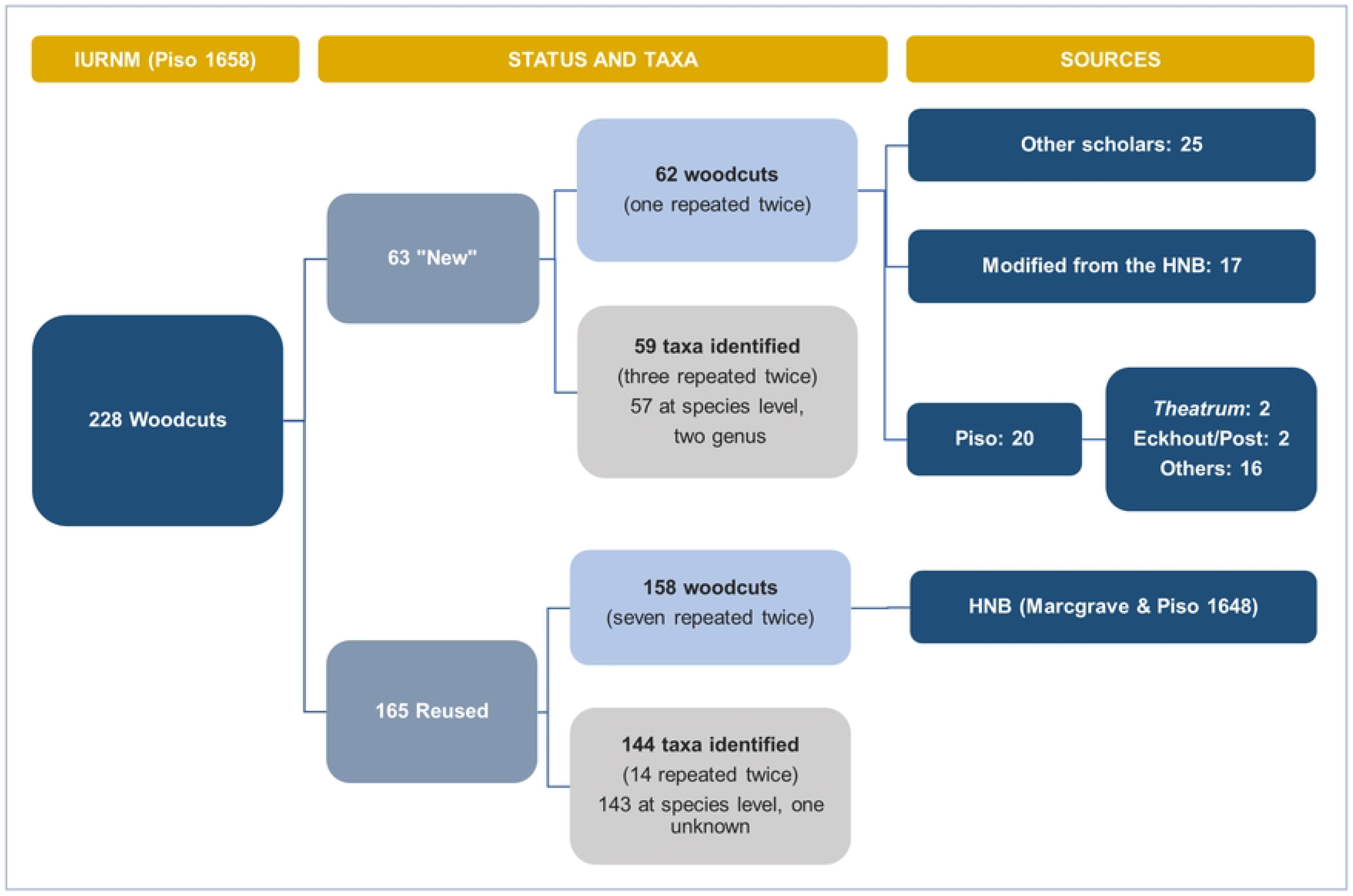
Tracing the sources of the plant woodcuts in the IURNM. Flowchart showing the organization of the woodcut images in the IURNM, the number of plant taxa, and the sources used to create these woodcuts.

Most of the woodcuts (72%) look exactly like those printed before in the HNB, suggesting these were made with the same woodblocks that De Laet had used ten years earlier, and which were kept by the Elzevier publishing house after his death (4). The remaining woodcuts (28%) were new illustrations used by Piso for the IURNM. Most of those new images (40%) were copied after Renaissance and Early Modern botanists and physicians’ herbals, such as those written by Brunfels (46), Fuchs (47), Matthioli (48), and Monardes (34,49) (S3, S4). Several images were “borrowed” from the works of Dodoens (50,51), De l’ Obel (52,53), and Clusius (35–37). Sometimes, Piso did not depict plants collected or observed in Brazil. To illustrate the neotropical *Gossypium barbadense* L. and *Astrocaryum vulgare* Mart., for instance, he copied the images of the cotton (*Gossypium arboreum* L.) and the African date palm (*Phoenix dactylifera* L.) from the work of Prospero Alpini (54,55), a Venetian physician who depicted the flora he encountered during his expedition in Egypt.

Several of those new woodcuts (28%) are modified copies made after the images in the HNB related to the same plant. Most of these modifications consisted in attaching the branch that was depicted in the HNB to a trunk – a style often encountered in late Renaissance herbals, likely to depict the tree habit of the plant while still showing enough detail of the fertile branch. Occasionally, some new images were created by combining parts of different plant species, something that Pickel (6: 23) defined as “fantasy woodcuts” (S3, S4: 29, 85, 87, 103). A few times, the modified plant parts in the woodcut were those that had nutritional or medicinal value. For example, *Cissampelos glaberrima* St. Hil. (*Caapeba*) is represented in the IURNM by two woodcuts (Fig 9a). One of them is the same as in the HNB (Fig 9b) and was made after one of the woodblocks that remained with Elzevier’s publishers. The other woodcut is new, albeit slightly similar to the first one but with larger roots that split in two (Fig 9a). Piso (44: 261) directed the reader to these different roots by citing this figure, and specifying that one of them became larger when it grew older. In addition, he experimented with the leaves and roots of *C. glaberrima* and documented its medicinal properties (44).

**Fig 9.**
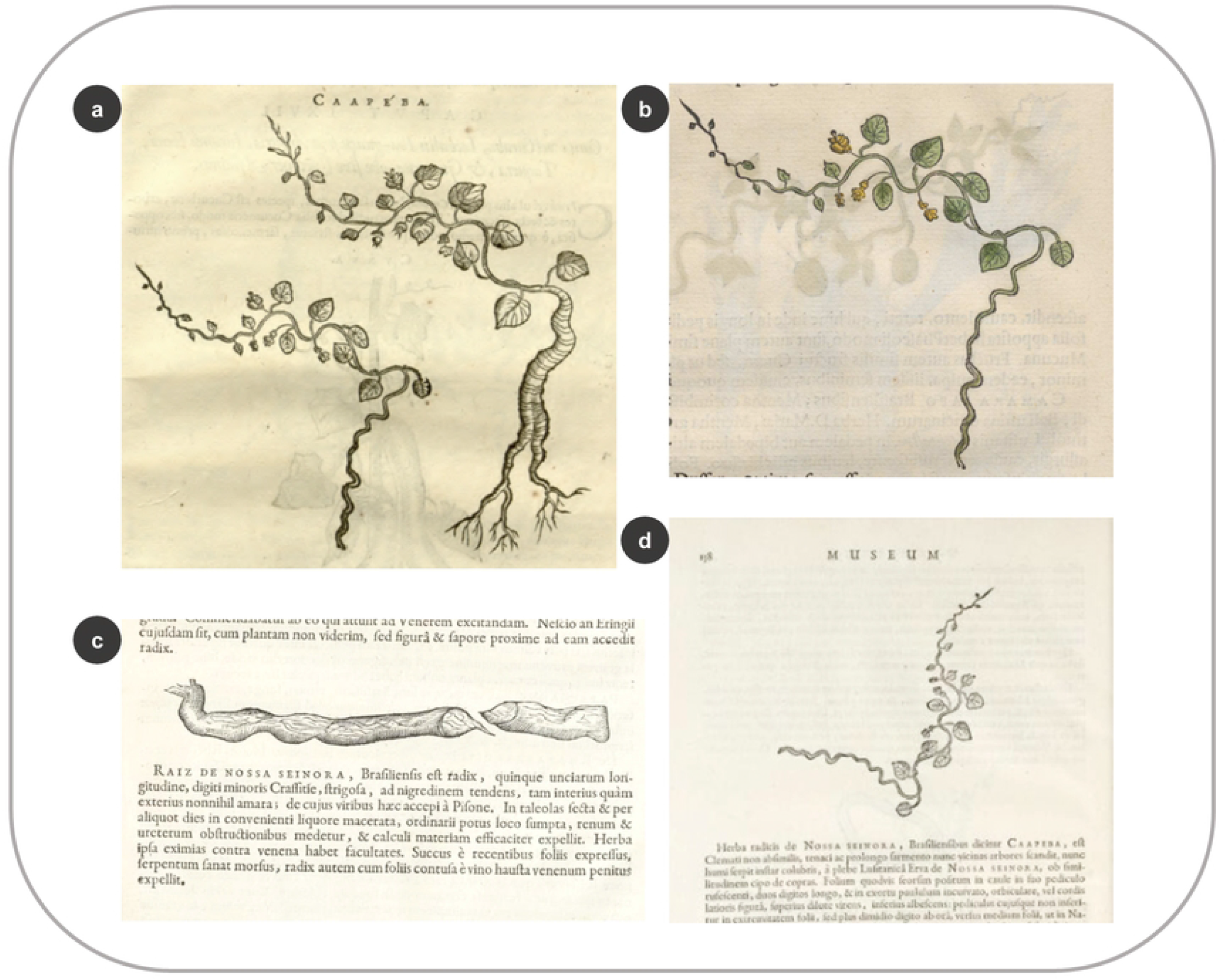
New image used by Piso (1658) and its connection to previously published woodcuts. (a) Woodcuts of *Cissampelos glaberrima* St. Hil. In the IURNM (page 261) (b) The same species in the HNB, documented by Marcgrave (p. 26) (c) Root of *C. glaberrima* shipped from Brazil as portrayed by Worm (1655: 157) (d) The same species made after the same woodblock (Worm 1655: 158).

Three years before the publication of the IURNM, Danish physician and historian Ole Worm published an image of *C. glaberrima* root in his book *Museum Wormiamum* (56). Worm obtained this root from Brazil (Fig 9c), perhaps as part of the plant material exchanged between De Laet and him (8). He also added the woodcut from Marcgrave by using the same woodblock used for the HNB (Fig 9d), which was owned by their common publisher: Elzevier (4,8). Just like Worm, Piso was aware of the medical relevance of *C. glaberrima* and emphasized its roots in his new image.

The remaining new woodcuts (32%) were not copied after the HNB woodcuts or images from other treatises. Four of them bear resemblances to the illustrations in the *Theatrum* and a few plants depicted by Eckhout or Post in their paintings (S3, S4: 1, 23, 39, 57). Most of them (15 woodcuts), we could not trace to any existent source. Piso could have had sketches and seeds of the plants on those images, but their exact origin or provenance remains uncertain. The physician did not provide the woodcut image of *Cuipouna* (*Cestrum schlechtendalii* G.Don,) because the drawing ‘had been damaged by the action of time’ (44: 178). He also lost the drawing of *Tapirapecu* (*Elephantopus mollis* Kunth) because of the ‘eventualities of the journey’ (44: 182), which may refer to events that occurred during his expeditions in Brazil, but also during or after his return to the Low Countries. Out of the 143 species preserved in Marcgrave’s herbarium, 55 are represented by a woodcut image in the IURNM, but only seven species appear in the newly made woodcuts and do not appear in the HNB. There is no resemblance between these seven images in the new woodcuts in the IURNM and Marcgrave’s herbarium vouchers.

## 3.3 Early modern ‘Photoshop’

### 3.3.1 Combining multiple sources

Grouping images from various sources was a technique that allowed us to portray a plant as completely as possible, especially when it was difficult to capture at once the different parts of the plant or when some details were missing. The medicinal shrub *Jatropha curcas* L. is represented by a woodcut in the HNB (Fig 10a) of lesser quality and scientific detail than its homolog in the *Theatrum* (Fig 10b). Despite the poor quality, De Laet included this image instead of making a new woodcut after the *Theatrum* oil painting. He either did not see this painting or he deliberately chose the drawing of lesser quality to avoid making a new woodcut design, thus saving time and money. The HNB woodcut was presumably made after a drawing by Marcgrave (6: 129). Ten years later, Piso published a different image, in which the internodal scars on the stem are visible (Fig 10c). This new image is slightly similar to the one in the *Theatrum*, although the ring-like scars on the stem are not visible in the oil painting. Marcgrave, however, documented this feature in p. 96 and compared it to a fig tree, which might explain why Piso added it to the image. Additionally, the seeds of *J. curcas* are displayed in both the HNB and the IURNM (Figs 10c, 10d). This image was made ‘au naturel’ after the seeds that De Laet (41: 137) received from Brazil and used in his treatise on the Americas (Fig 10e).

**Fig 10.**
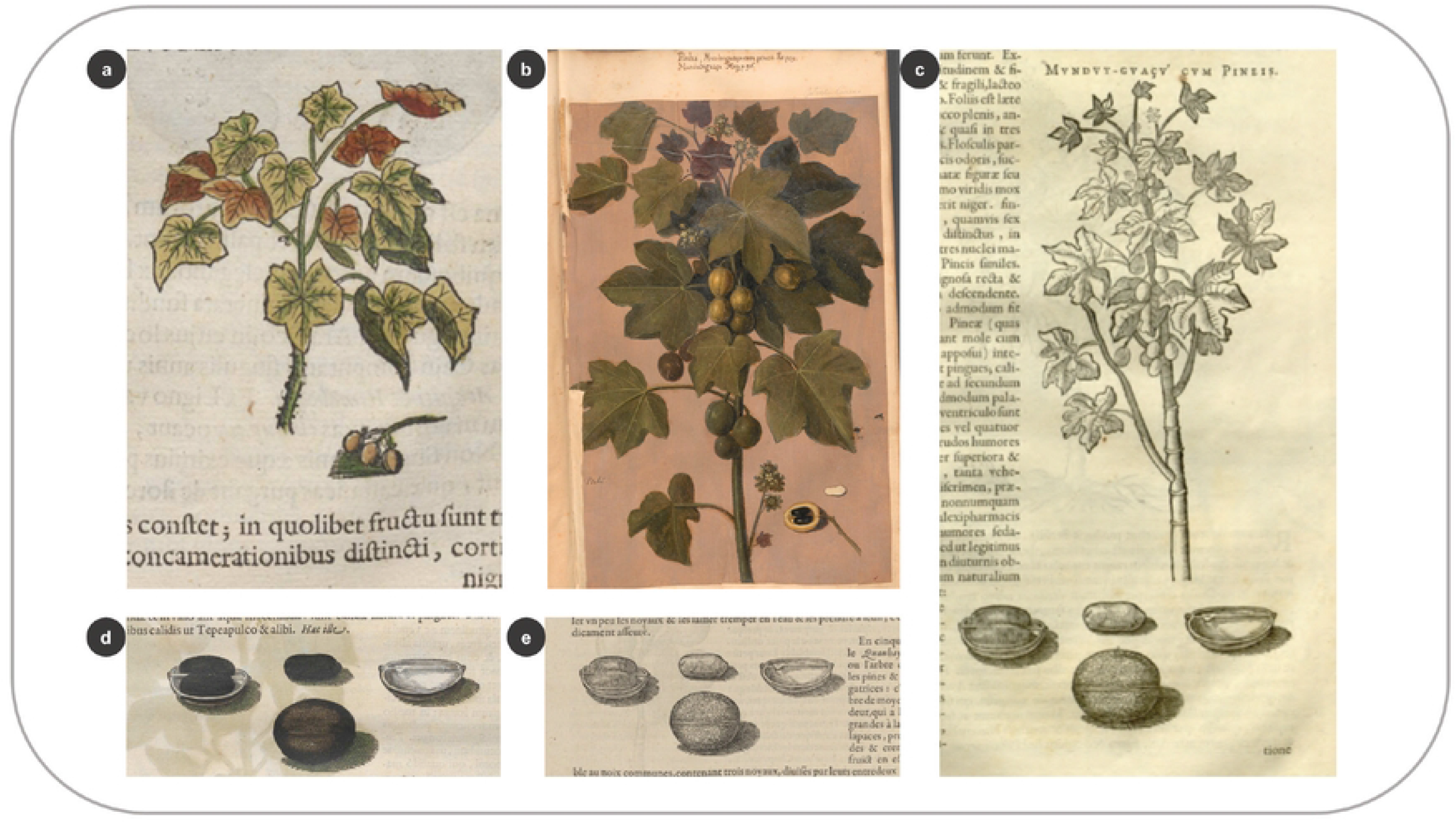
Combining multiple sources to create the woodcuts for the natural history books. (a) Woodcut of *Jatropha curcas* L. in the HNB (Marcgrave 1648: 96) (b) Oil painting of the same species in the *Theatrum* (f. 199) (c) Woodcut of this plant and its seeds in Piso (1658: 179) (d) Woodcut of *J. curcas* seeds in the HNB (Marcgrave 1648: 97) (e) Original woodcut of the seeds (De Laet 1640: 136).

Another example of the use of multiple sources occurs with *Bixa orellana* L., known today as *Achiote, Annatto*, or *Urucu(m)* in Brazil – among others ([Dataplamt](http://www.dataplamt.org.br/v3-novaversao-block/#/listagem/?nomecientifico=bixa%20orellana), accessed 23.09.22). The woodcut in the HNB shows a flowering branch with four terminal fruits and small fruits on a lateral branch (Fig 11a). The flowers are missing in the *Theatrum* illustration, in which some of the fruits are depicted as open and full of red seeds (Fig 11b). These seeds were one of the earliest trade goods exchanged between Indigenous peoples and Europeans in South America and were exported to Europe in the mid-sixteenth century to be used as a dye, colorant, and cosmetic (57,58). The two images differ greatly, and although the fruits of *B. orellana* were of great economic relevance, De Laet included the woodcut with the less showy and accurate image of them, possibly made after a drawing by Marcgrave. In the IURNM, the fruits of *B. orellana* are placed together with the HNB woodcut (Fig 11c). Piso borrowed the image of a fruiting branch from *Exoticorum Libri Decem* by Clusius (36: 74) (Fig 11d). He added an image of an open fruit with seeds, which curiously looks as if the fruit on the left of the branch had been taken, opened in half, and laid on the bottom of the figure (Fig 11c). Clusius obtained the original branch from the aristocrat and naturalia collector Pieter Garet (c.1552/5-1631), who wrote to the botanist that Brazilian Indigenous peoples used the seeds to color their bodies red (59).

**Fig 11.**
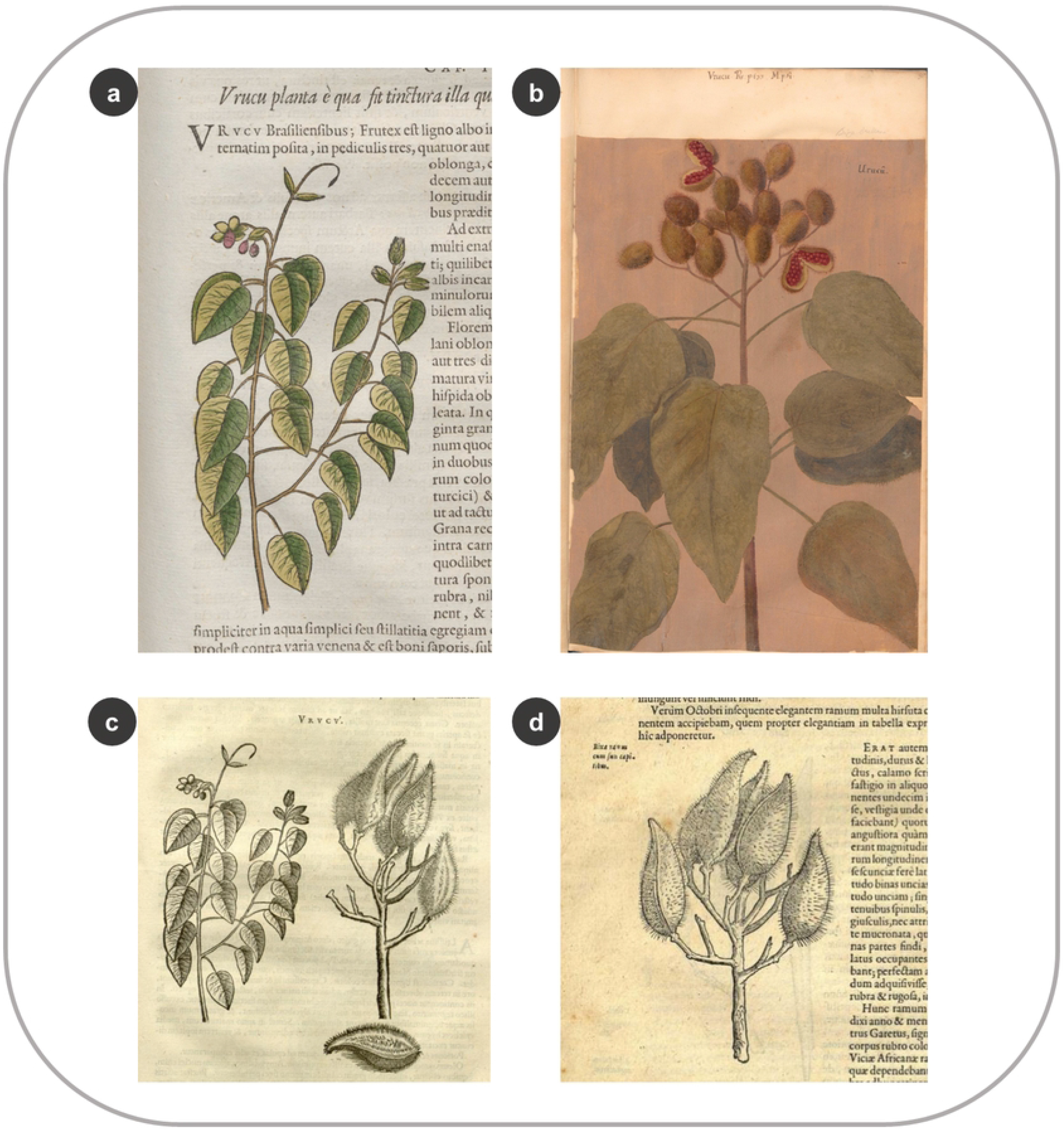
Use of multiple sources to create the plant woodcuts by De Laet and Piso. (a) Woodcut of *Bixa orellana* L. in the HNB (Marcgrave 1648: 61) (b) Oil-based illustration of *B. orellana* in the *Theatrum* (f. 95) (c) Woodcuts of the same species in the IURNM (Piso 1658: 133) (d) *B. orellan*a branch in Clusius’ treatise (1605: 74).

### 3.3.2 Combining different stages of plant life

Several woodcuts show flowering and fruiting stages depicted together. In the HNB, the woodcut of *Crateva tapia* L. bears a fruiting branch, which includes an open fruit full of seeds (Fig 12a). In the same woodcut, we also see the inflorescence and the tiny (immature) fruits, characteristic of the Capparaceae species (Fig. 12a). *C. tapia* in the *Theatrum*, however, is represented by two illustrations glued in the same folio and each depicts a fruiting and a flowering branch (including the fruit buds) (Fig 12b).

**Fig 12.**
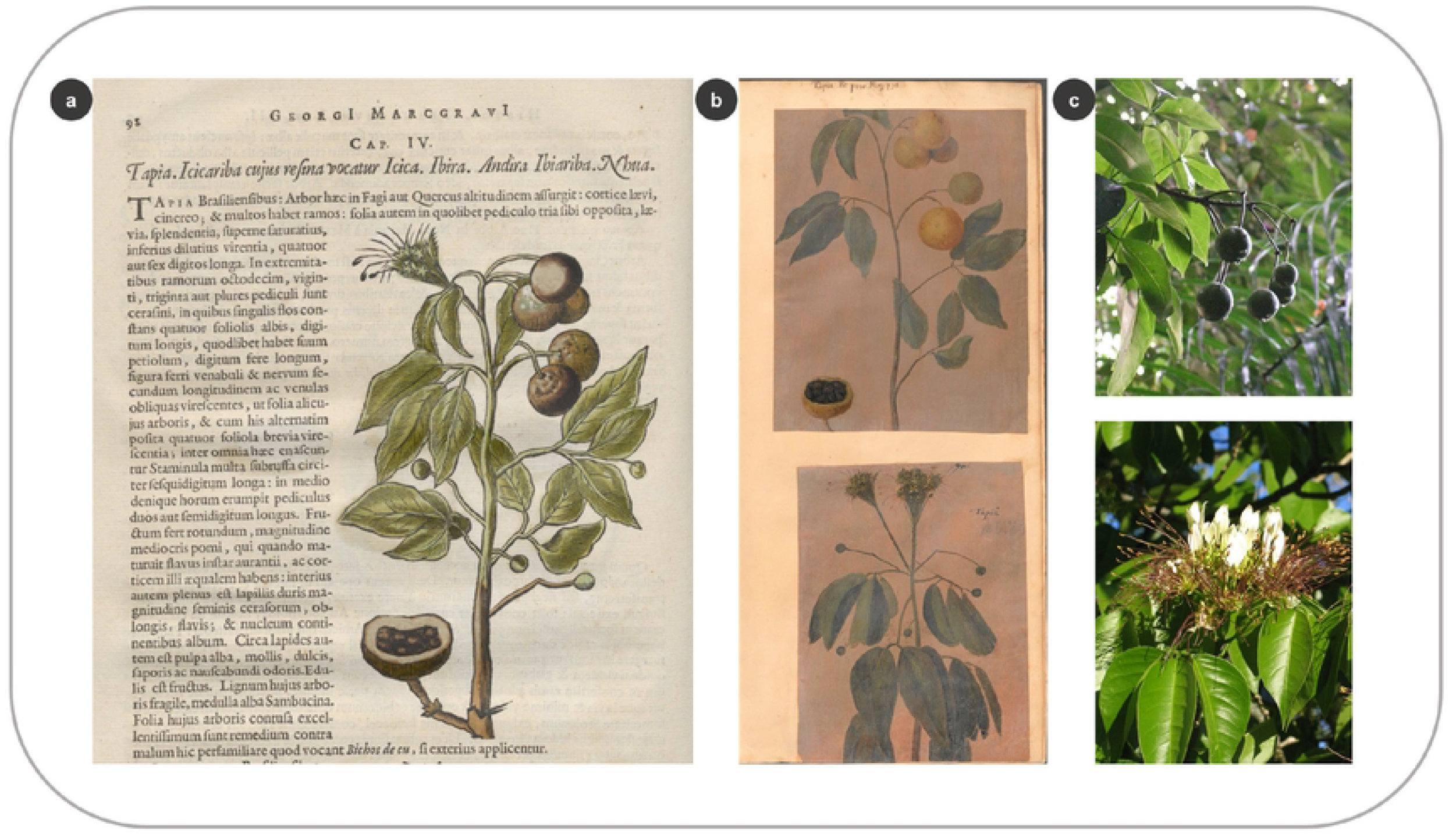
Combining different stages of plant life in one image. (a) Woodcut of *Crateva tapia* L. in the HNB (Marcgrave 1648: 98) (b) Fruiting branch of the same species in the *Theatrum* (above) and flowering branch (below) (f. 113) (c) Fruiting branch of *C. tapia* (above), and flowering branch (below) (by T. Leão and A.S. Farias-Castro, in: www.flickr.com).

Flowering and fruiting stages of *C. tapia* can overlap in nature (60) (example in Species Link: HUEFS 134255, identified and collected by Lyra-Lemos R.P. Alagoas, Brasil) but this is not always the case (Fig 12c). However, the oil paintings show the different fertile structures separately, which must have belonged to branches with different reproductive stages or collected at different times. Either way, the image depicted in the HNB – similar to those two separate illustrations – conveyed more botanical information in one image and saved money during the printing process. After all, preparing and including images in treatises was an expensive endeavor in the early modern period (61,62), as it is today.

Several images merged flowering and fruiting stages, both in the *Theatrum* and the HNB, such as *Annona montana* Macfad. and *Byrsonima cydoniifolia* A. Juss (S2: 331, 411), of which the reproductive stages can indeed appear simultaneously in nature (https://specieslink.net/search/). Hence, we cannot infer whether these images were based on separate flowering and fruiting individuals– as was the case for *C. tapia*. We noticed, though, that these species were drawn with a larger number of leaves in the *Theatrum* than in the HNB. In contrast, other images are represented the other way around: *Paullinia pinnata* L. in the *Theatrum* (f. 283), whose infructescence and open fruit resemble the woodcut in the HNB (32: 114), shows many more (and slightly different) leaves (S2: 61).

## 3.4 Connections between De Laet’s manuscript and the woodcuts

Our database that connects the images and descriptions in the manuscript to their species descriptions and woodcuts in the HNB is provided in Supporting information S5. In total, we counted 388 plant descriptions and 165 plant entries with the word *Icon* next to the entry (S5). Out of these entries, 20 have written ‘in lib. [x number]’ (*in lib* meaning ‘in the book’). The entry on *Mureci* (*Byrsonima cydoniifolia* A.Juss.) lacks the word *Icon*, but it has ‘in lib. 93’ written next to it, thus it was likely associated with a specific drawing kept in a notebook that was numbered 93. Out of these 166 entries (*Icon* ones plus the entry on *Mureci*), we identified 164 species; two of them remained unidentified as we could not match them to the descriptions in the HNB (S5). There are 149 *Icon* entries (including *Mureci*) whose plant descriptions include a woodcut in the HNB. Surprisingly, we found 15 entries that include the word *Icon*, but the species that are described in those entries are not depicted by a woodcut in the HNB. The drawings could have existed, but they were probably misplaced by De Laet. For instance, *Tapyracoaynana* (*Cassia grandis* L.f.) in De Laet’s manuscript bears the word *Icon*, but it has no image in the HNB. The woodcut of *Byrsonima sericea* DC. is depicted near its description (3: 134) and this was probably the corresponding image tagged with *Icon* in the manuscript (instead of *C. grandis*). This also confused Piso who, ten years later, placed the image of *B. sericea* with a woodcut of a *C. grandis* pod and called both *Tapyracoaynana* (44: 158).

Noteworthy are the plant entries of this manuscript and their correlation to the *Theatrum* (S5). Most species (109 spp., 66%) tagged with *Icon* (thus mostly have a woodcut in the HNB) are not present in the *Theatrum*. The remaining (57 spp., 34%) have a correspondent species within the oil paintings, and around half of their woodcuts (27 spp.) are similar to the *Theatrum* (S2), with a few of them (8 spp.) with ‘in lib. [x number]’ written (S5). We also found this addition in 13 of the *Icon* entries, but the corresponding woodcuts are not similar to the *Theatrum* images. One of the entries reads *Icon in p. 172* and corresponds to *Furcraea hexapetala* (Jacq.) Urb., an image that was used by De Laet in his treatise on the Americas. This number does not match the page number for the same image/species in the editions we reviewed (40: 666, 41: 608, S2: 321-322).

By studying how the images in De Laet’s manuscript are arranged, we observed that nine drawings in the *verso* folios do not bear the word *Icon* in their descriptions in the *recto*, but they have similar names. Out of those, eight are represented with an identical woodcut in the HNB (4,8), of which five are only in Piso (32). This is strange, as De Laet presumably used this text to elaborate on Marcgrave’s chapters on plants. Eight drawings (four pencil-based and four proof woodcuts) resemble the *Theatrum* illustrations (non-reversed). The drawings were glued to the *verso* folios in De Laet’s manuscript after the completion of the manuscript (perhaps by another person) by matching the vernacular names that were written in the recto with the plant descriptions. An exception is *Lecythis pisonis* L., which lead drawing lacks a name and was placed somewhere else (S5).

## 3.5 Connections between the images and their vernacular names

All the HNB woodcuts that look similar to their correspondent taxa in the *Theatrum* share the same or similar (cognates) vernacular names (S1). All the entries with the remark *Icon in Lib x* in the manuscript (n=8) are associated with plant taxa that bear similar images and names in both sources (S5). There are some peculiarities to it: *Camara uuba* in De Laet’s manuscript bears *Icon in lib. 69* next to it (Fig 13a). Therefore, we would assume that the described plant would include a woodcut in the HNB copied from a book or notebook numbered 69. The description corresponds to *Calea elongata* Baker (6), but the woodcut in the HNB is very similar (in reversed format) to the *Theatrum* illustration for *Lantana camara* L. (Fig 13b). The oil painting bears the name *Camaràuna* (Fig 13c), which was presumably written by the artist in question (Fig 13d). De Laet warned the reader in the HNB that ‘the image we give here [with *C. elongata* description], even though we found it under the name *Camara uuba*, seems to be from another *Camara*, which the author [Marcgrave] mentioned before’ (De Laet’s commentaries in (3: 6) (Fig 13e).

**Fig 13.**
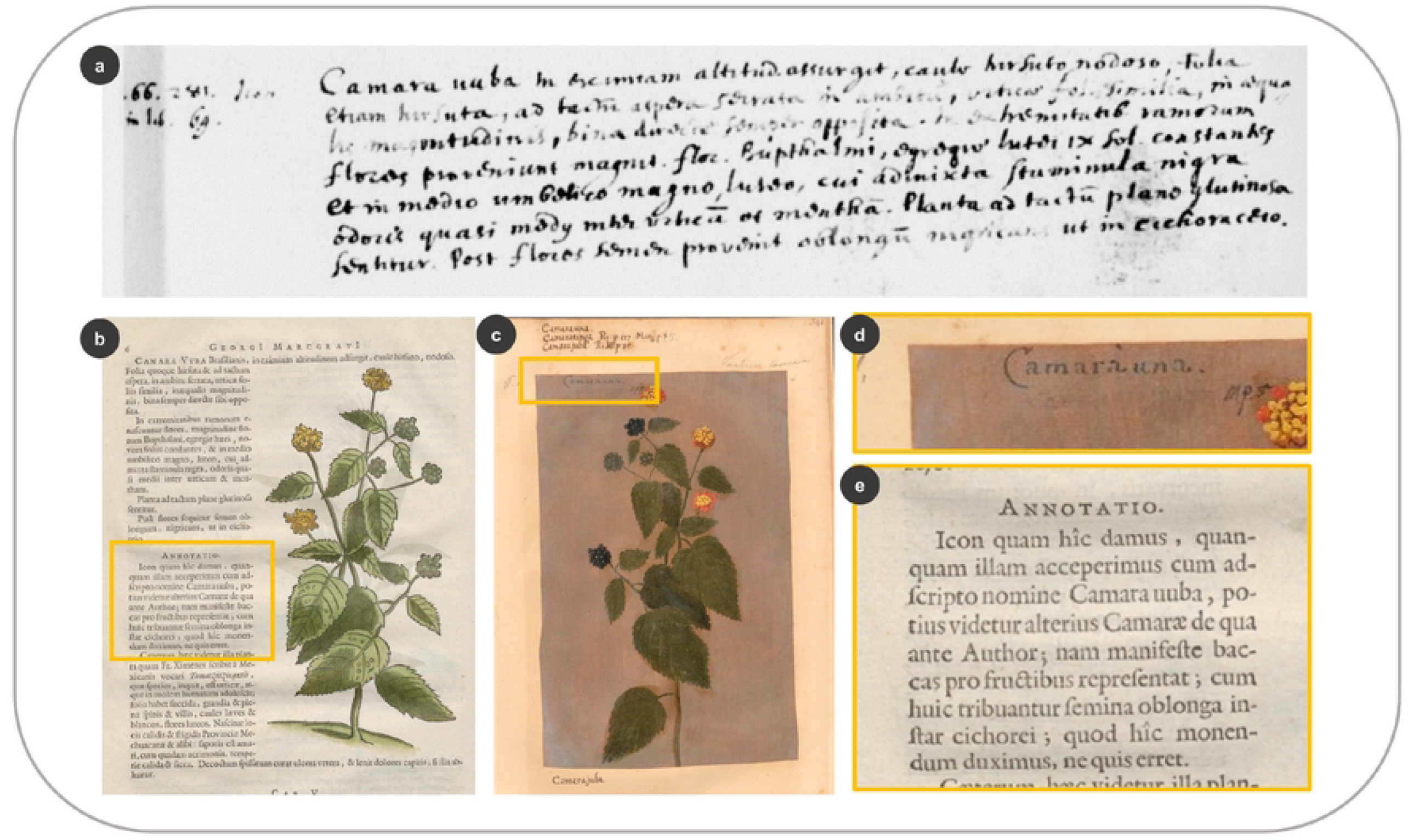
Linking the woodcuts in the HNB with De Laet’s manuscript and the *Theatrum*. Description of *Calea elongata* Baker next to *Icon in lib. 69* in De Laet’s manuscript (f. 5r) Woodcut of *Lantana camara* L. in the HNB (Marcgrave 1648: 6) (c) Oil painting of *L. camara* in the *Theatrum* (f. 341) (d) Vernacular name written on the illustration (e) De Laet’s annotation on the use of this image (Marcgrave 1648: 6).

Most of the taxa that have different vernacular names in the HNB and the *Theatrum* are also illustrated differently, except for three species. The woodcuts and illustrations of *Spighelia anthelmia* and *Dorstenia brasiliensis* Lam. look alike, but do not bear any name in the oil paintings. The names could have been cut out, as happened for some of the oil paintings before being glued into the bound collection (11: 275). *Samanea saman* (Jacq.) Merr. is named *Guaibí pocaca biba* in the HNB (3: 111) and *Nhuatiunana* in the *Theatrum* (f. 399). Since their images are only slightly similar, this weakens the possibility that they originated from the same source.

Another species that has different local names in the HNB and the *Theatrum*, and is represented with a different image, is *Schinus terebinthifolia* Raddi. A fruiting branch in the HNB (Fig. 14a) is accompanied by the description of this species and the names *Aroeira* and *Lentiscus*. The woodcut, however, belongs to another taxon. De Laet reused the figure (Fig 14b) from his book on the Americas, in which he described a Peruvian tree (41: 327). He copied this image after the treatise *Curae Posteriores* (37: 94), in which Clusius commented on the work of the Spanish physician Monardes on American plants (49: 39). This woodcut represents *Schinus molle* L., not a Brazilian species, of which plant material and knowledge were circulating in Clusius’ network of physicians and naturalia collectors since the beginning of the seventeenth century (63).

**Fig 14.**
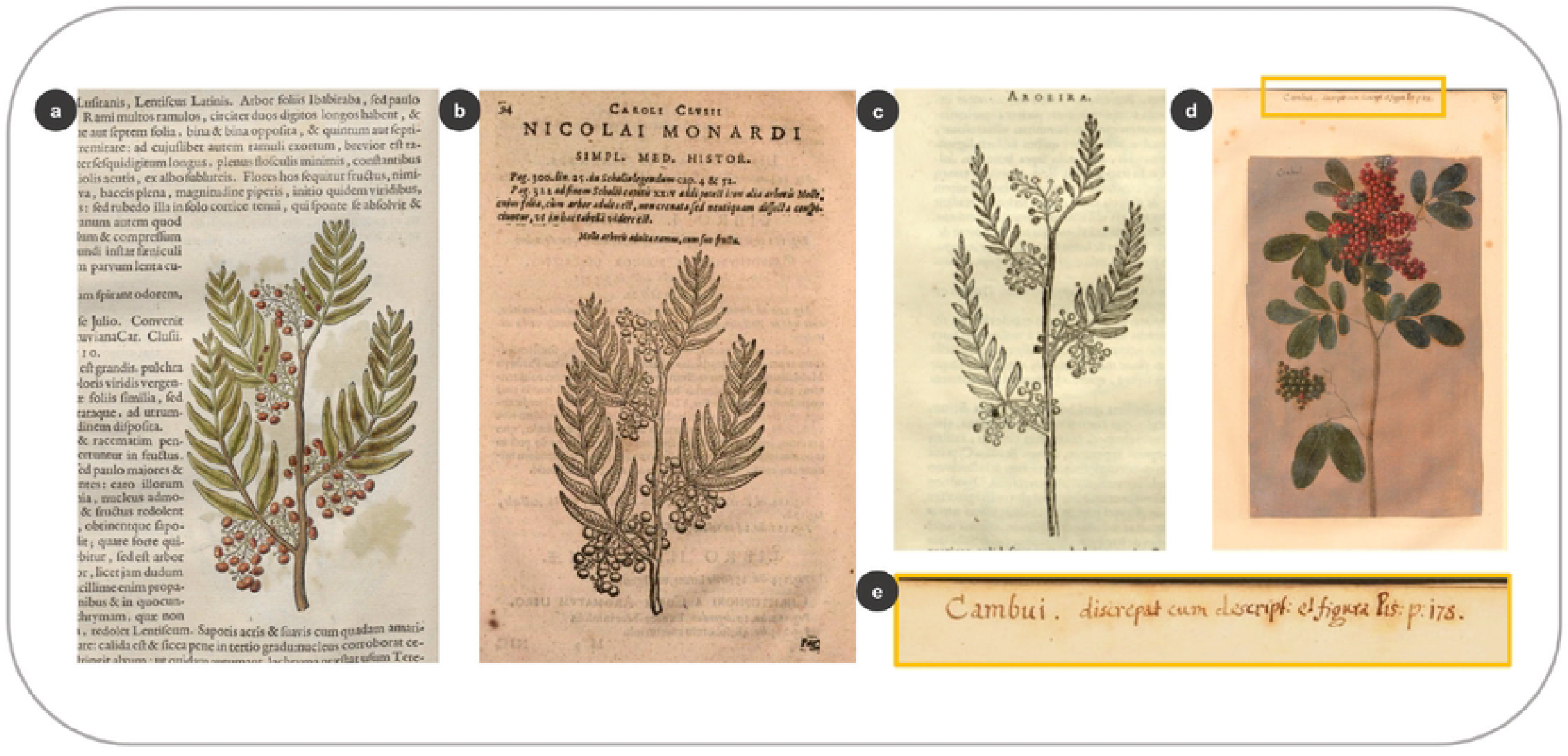
Different vernacular plant names for the same species resulted in mismatching images and descriptions. (a) Woodcut of *Schinus molle* L. (*Aroeira*) in the HNB with a description of *S. terebinthifolia* Raddi. (Marcgrave 1648: 90) (b) the same woodcut of *S. molle* in Clusius (1611: 94) (c) Similar woodcut of *S. molle* in Piso (1658: 132) (d) Illustration of *S. terebinthifolia* in the *Theatrum* (f. 295) (e) Close-up of the annotation in the *Theatrum* (f. 295) which includes the name *Cambuí* and the misleading reference to Piso (1658: 178).

Ten years later, the woodblock must have been damaged or disappeared, as Piso (2) used a slightly different image based again on De Laet’s and Clusius’ woodcuts (Figs 14b, 14c), instead of the legitimate species, as the one present in the *Theatrum* (Fig 14d). This oil painting includes the name *Cambuí* (Fig 14d), which we found in the HNB, but for a different species: *Eugenia involucrata* DC., in (32: 82). Whoever wrote the reference above the illustration of *S. terebinthifolia* in the *Theatrum* was pointing out the woodcut of *E. involucrata*, but they noticed the discrepancy between the image and the description (Fig 14d).

The names *Cambuí* and *Aroeira* were used for *S. terebinthifolia* in seventeenth century Dutch Brazil and are used for the same plant today ([Dataplamt](http://www.dataplamt.org.br/v3-novaversao-block/#/listagem/?nomecientifico=schinus%20terebinthifolia), accessed 23.09.22). The name *Cambuí/i* is currently used for several species within the Anacardiaceae, Myrtaceae, and Fabaceae families ([Dataplamt](http://www.dataplamt.org.br/v3-novaversao-block/#/listagem/?nome=cambuy), accessed 23.09.22). Hence, if the names are different for the same species, most of the images do not bear strong resemblances. Nevertheless, the same or similar names can be associated with different images for the same taxa, as happens with *Bixa orellana*, named *Urucu* in both the HNB and the oil painting (Fig 11). Interestingly, most of the vernaculars in the oil paintings are in an Indigenous, Tupi-based language, and just a few originated from an African language. This contrasts with the HNB, which apart from the Tupi names, often provides Portuguese, and to a lesser extent Dutch, Spanish, and Latin plant names (64).

## 4 Discussion

### 4.1 ‘Call me by your name’: organizing the Brazilian flora by their Tupi names

Arranging the visual and textual material to create the natural history treatises was an arduous task. As the systematic binomial nomenclature had not yet been established, De Laet, Mentzel, and other scholars who used the HNB as a reference for tropical plants, such as C.F.P. von Martius, struggled to organize the megadiverse Brazilian flora by their vernacular names (65). They cross-referenced the different sources that were available to them, just as we did in this study – although we used the scientific names first.

The difficulties of associating the visual material with the right vernaculars and their corresponding descriptions must have contributed to the development of taxonomy in combination with the flow of exotic plants (either dried, alive, or illustrated) to botanical gardens and private collectors (66,67). Linnaeus used the descriptions of Marcgrave and Piso in his *Systema Naturae* (68), and he often used the Tupi Indigenous names as generics or specific epithets (19) – like for *Crateva tapia*, in which *tapia* is a Tupi-based vernacular reported by Marcgrave.

That most of the images are labeled with Tupi-based and a few African plant names imply that artists, naturalists, and Indigenous and enslaved Africans had worked in close connection with each other, as it occurred for the creation of the textual sources in the HNB (30). Piso stated in his preface that he ‘submitted to the examination and practice everything that out of the vast theater of nature [he] observed or received by the Indigenous veterans’ (2: 6, 43: 8). Moreover, the painter traveling with Piso into the interior could have been an enslaved African. All members of Johan Maurits’s retinue were entitled to personal assistants, as is stated in a document from the WIC at the National Archives in Den Haag (69). To what extent those contributions were forced upon the local inhabitants is not explicitly mentioned in the archives or natural history books. Nevertheless, such connections were certainly uneven – and likely unjust – due to the structural violence that prevailed in the colony. The military activities of the WIC, which ensured the company and their stakeholder’s economic profitability overseas, were sometimes combined with scientific expeditions (70). Slavery and armed operatives to subjugate local peoples who did not ally with the WIC were part of these activities (71,72) Our database with all species listed in the HNB and the IURNM, Marcgrave’s herbarium, and the *Libri Picturati* together with their vernacular names (S1, S2, S3, S4) provides a good ground to analyze western and non-western plant nomenclature systems and their preservation over time, as well as to review the role of Indigenous and African knowledge embedded in these natural history collections.

### 4.2 Visual knowledge-making processes in the HNB and the IURNM

By analyzing the plant woodcuts, we can observe that a variety of models were used to represent the different plant species: freshly picked plants, living individuals, dried fruits, seeds, branches, or herbs, sometimes preserved as herbarium vouchers. Not all images originated in the immediate surroundings of Johan Maurits’ palace (16,24), as plants were gathered in a variety of places and by different people, on several occasions by the Indigenous Brazilians. To portray all fertile stages of a plant was important, but sometimes this aim failed due to inconveniences during the expeditions or the journey back to Europe, or because seeds failed to germinate or grow into plants in Holland. Exsiccates were crucial in the production of botanical books as they allowed authors to compare the specimens with the engravings (73), although only a fraction of the woodcuts in the HNB were based on the vouchers collected by Marcgrave. Unlike the Renaissance authors of popular herbals such as Fuchs or Gessner (1516-1565), who remained in control of the publishing of their work and could correct botanical mistakes (74,75), Marcgrave could not intervene to review his draft chapters or his published work because of his premature death. Instead, De Laet, who had never crossed the Atlantic and did not have botanical training, took the lead in assembling the notes and images that were given to him by Johan Maurits to create the HNB.

This material originated from a variety of people, who specialized in different subjects, but whose skills were connected. Marcgrave was trained as a botanist and astronomer, but he also made drawings and retrieved medicinal plant knowledge from the native population, just as the physician Piso did. Marcgrave, however, was not only interested in the utilitarian value of the flora he studied, hence some of his woodcuts belong to species that were not described as useful in the HNB. These images were not represented in the IURNM (S4) because Piso aimed to create a more pragmatic field guide. Piso stated that he created his book ‘with engravings copied after nature [*iconibus ad vivum depictis*], not only for the delight and admiration of the reader but, above all, to serve the doctors and the sick’ (2: 47). This statement is partly true: his images were not original *sensu strictu* because he mostly reused the same woodblocks that were used for the HNB, but the majority of them originated from the floristic studies in Brazil by his colleague Marcgrave.

When sources were available, De Laet and Piso commissioned figures that were carved onto the woodblocks; often by looking at the vernacular names that accompanied those sources, as we observed for the *Theatrum*, in which the overlapping images with the HNB shared the same vernacular names. Then, they ordered the design of a plant drawing to be further transferred into a woodblock, only if other images or specimens were lacking. Several accurate botanical representations included in the oil paintings, whose names matched those in the HNB, were not carved onto the woodblocks. Other images must have been already available, which ended up being the basis for the woodcuts. This strategy allowed them to economize time and money. However, Piso added several modifications in the IURNM, especially for the trees, so he altered the woodblocks or had new ones made based on the HNB images.

Following Chen (76), we argue that the HNB created a new visual language by including many images never seen before. Those images constituted a legitimate visual repertoire that was later borrowed by others, such as Worm (56) or Piso (2), whose IURNM resulted in an accumulation, rather than an innovation, of visual knowledge by replicating the images published in the HNB a decade before. Piso also copied the woodcuts from other authors, especially those used by the Plantin publisher’s house in the sixteenth and seventeenth centuries (S4), even though these plant images were often based on different locations. He sometimes even used images of Mediterranean, Asian and African species that did not occur in Brazil at that time or were not reported in the HNB.

### 4.3 Provenance of sources for the plant woodcuts

The question that Whitehead and Boeseman (4) posed on what extent the *Libri Picturati* images served as models to elaborate the woodcuts of the HNB is answered in this research. The *Libri Principi*’s plant images were not used as the basis for any of the woodcuts in Marcgrave and Piso’s treatises. This differs from the animal woodcuts in the HNB, which bear a strong resemblance to the watercolors in the *Libri Principis* (12,15,19) The *Theatrum* had some influence, but it did not constitute the main basis for the plant woodcuts (4,16), as only one-third of the woodcuts resemble the oil paintings. The majority of the woodcuts were based on other sources, most likely on the drawings made by Marcgrave in Brazil. The naturalist and his assistants could be acknowledged as the author(s) of most of the flora woodcuts, instead of Eckhout (15). Despite a few drawings embedded in De Laet’s manuscript, whose authorship has been attributed to Marcgrave (4,11), there are no other records of the naturalist’s original drawings made in Brazil, and even so, this authorship cannot be fully attributed to Marcgrave because some bear a strong resemblance to the *Theatrum*’s illustrations. Hence, if Eckhout made these oil paintings, then the agents behind most of the woodcuts would be primarily Marcgrave and to a lesser extent Eckhout, as indicated by Brienen (16). There are 163 botanical woodcuts in the HNB (from Piso and Marcgrave’s chapters on plants) that cannot be traced to the *Libri Picturati*, Marcgrave’s herbarium vouchers used by De Laet, or other scholars’ works. This number represents nearly half (47%) of the drawings that Marcgrave mentioned he had made in his letter to De Laet in 1640 (28). Considering that the naturalist kept working in Brazil until 1643, a larger number of drawings can be expected, several of which were lost or never ended up in De Laet’s hands. Another possibility is that Marcgrave was the author of the models that were used to elaborate the oil paintings or at least some of them. Apart from some of the drawings glued in the *verso*, several entries (27 species) in De Laet’s manuscript with the word *Icon* correspond to the oil paintings images. These could correspond to previous drawings of the oil paintings included in a numbered notebook created in Brazil (15). As happened with the animal woodcuts and the *Libri Principis*, some plant woodcuts show greater details than their corresponding images in the *Theatrum*. Others, though, show a less crowded image (e.g., by reducing the number of leaves) than their overlapping images in the *Theatrum*. The existence of plant studies previous to the oil paintings, which were later used as models to create the *Theatrum* ones (15), is reasonable. The increased value of the Brazilian imagery lies in the fact that many illustrations and drawings were produced *in situ* (16). Nevertheless, the oil paintings could have been made in the Dutch Republic, as similarly hypothesized by Johann Horkel (1763-1846) for the watercolors of the *Libri Principis* (25). After all, this iconographic material was not given to the Elector of Brandenburg until 1652, eight years after Maurits’s return to the Low Countries.

### 4.4 Future research and recommendations

Further studies of the fauna woodcuts in the HNB and IURM and their correlations with the animal illustrations in the *Libri Picturari* are crucial to complement existing studies (4,5,15–17,20,21,23,26), to compare our research outcomes and shed more light into the sources used for the woodcuts. Yet, the location of many sources is unknown or even lost. To solve these mysteries, archival research should be conducted alongside the study of herbaria, libraries, and private collections that are linked to material originating from Dutch Brazil. Digitizing the known sources is essential to facilitate their study without the need of touching the fragile material. Publication of the high-resolution images in an online open format would increase its access to a larger academic community. However, this does not guarantee its dissemination to the wider public. It is pertinent to work towards collaborative projects between Indigenous and Afro-Brazilian communities, researchers, and representatives of the western institutions that hold this biocultural material. Recently, various historical Brazilian materials have been used in successful cross-cultural projects that generated valuable outcomes for the Indigenous communities involved (77–79).

## 5. Conclusions

The repertoire of drawings that were used to elaborate the HNB and IURNM is incomplete. Creating these treatises can be compared to making a puzzle with several pieces lacking, and the impossibility of coming back to gather them. Nevertheless, our systematic analysis reveals new insights about the sources of the woodcuts in these books. The images embedded in these natural history books reflect an intentional effort towards portraying as much botanical information as possible. This goal was achieved by using local people’s knowledge of the environment to provide the plant material that was later captured in images and perhaps to assist further with the artistic process. Moreover, Tupi-based plant names played a key role in the arrangement of textual and visual sources, which were sometimes confusing for western scholars. Availability and economy were crucial, so most of the drawings that were carved onto the woodblocks were those that arrived from Brazil, sometimes combined with existing images of different provenance, and a few new ones made in the Dutch Republic.

Overall, the process of visual knowledge-making differs between the two books: the HNB mostly relied on primary visual sources to depict the flora, while the IURNM relied on secondary ones. Yet, the human agency behind such sources requires further attention. Archival studies and collaborative projects with Indigenous and Afro-Brazilian communities and researchers, art historians, and artists could shed light into the missing pieces of this conundrum and the multiple (hidden) histories related to these collections. There is certainly far more than botanical imagery behind the nature portrayed in Dutch Brazil; but to be able to see it, different eyes have to look at it.

## Acknowledgments

We greatly thank Jagiellonian library curator Izabela Korczyńska and the British Library (Manuscript Department) for making possible this research by providing us the digital images of the *Theatrum* and the scanned copies of De Laet’s manuscript. Likewise, we are grateful to information specialist Godard Tweehuysen for providing access to the HNB kept in Naturalis Biodiversity Center. Our gratitude goes to the plant taxonomists and ethnobotanists who reviewed the woodcuts and botanical illustrations and advised us with their expertise. Special thanks to Abisaí J. García Mendoza, Eduardo Hajdu, Jorinde Nuytinck, Nicole de Voogd, Thomas W. Kuyper, and Viviane S. da Fonseca Kruel. Many thanks to Huib Zuidervaart for sharing his exhaustive research on Marcgrave – or as he found out, Marggrafe. Lastly, big thanks to our colleagues Carolina Monteiro, for meticulously digging into the National Archives in Den Haag and sharing crucial data on slavery in Dutch Brazil, and Csilla Ariese for facilitating the use of the Filemaker software and hence contributing to the completion of the visual appendices.

## Supporting information captions

S1. **Sources of the plant woodcuts in the *Historia Naturalis Brasiliae* (Marcgrave and Piso 1648)**.

This table provides an overview of the plant taxa that were depicted in the HNB. To avoid repetitions, the woodcuts in common to both Marcgrave and Piso for the same species are only depicted in Marcgrave (1648). We analyzed the image’s overlaps between the HNB and older or contemporary sources that included botanical images of the same species.

Additionally, we added the vernacular plant names as documented in the HNB and the *Theatrum* illustrations to facilitate our comparative analysis between plant names. All the plant images can be visualized in the appendix (S2).

**S2. Nature portrayed in images in Dutch Brazil – Appendix (S2) Sources of the plant woodcuts in the *Historia Naturalis Brasiliae* (1648) Database**.

This PDF file includes the repertoire of woodcut images of the plants depicted in the HNB and their corresponding images retrieved from older or contemporary sources by cross-referencing their scientific names. This appendix allows us to visualize – among botanical illustrations, herbarium vouchers, plant sketches, and other plant materials that crossed the Atlantic – the potential sources that were used to elaborate the woodcuts in the HNB.

S3. **Sources of the plant woodcuts in the *India Utriusque re Naturale et Medica* (IURNM, Piso 1658)**. This table provides an overview of the plant taxa that was depicted by Piso (1658) with different woodcuts than in the *Historia Naturalis Brasiliae* (HNB, Marcgrave, and Piso 1648) and the presence of the same species in related sources. The similarities among visual sources allow us to hypothesize about the provenance of the woodcuts. These (dis-)similarities can be visualized in an appendix (S4). Correlations on the plant woodcuts in common with the HNB are analyzed in the main paper based on the dataset (S1), and the plant images are portrayed in a PDF file (Appendix S2).

S4. **Nature portrayed in images in Dutch Brazil – Appendix (S4) Sources of the plant woodcuts in the *India Utriusque re Naturale et Medica* (1658) Database**

This PDF file includes the repertoire of distinctive plant woodcuts included in the IURNM (i.e., those that did not appear in its precedent work, the HNB) and their corresponding images retrieved from older or contemporary sources by cross-referencing their scientific names. This appendix allows us to visualize – among botanical illustrations, herbarium vouchers, plant sketches, and other plant materials – the potential sources that were used to elaborate the woodcuts in the IURNM.

S5. **Correlations between the plant woodcuts in the HNB and the botanical annotations in De Laet’s manuscript**.

This table shows the correlation between the plants that presumably originated from a sketch or drawing that came from Brazil in a numbered booklet (marked with the word ‘Icon’ in the recto folios of the manuscript), the plant woodcuts in the HNB, and their corresponding images in other sources from Dutch Brazil (like the *Theatrum* and Marcgrave’s herbarium). The verso folios of the manuscript correspond to plant drawings or proof-woodcuts glued to these folios.

## References

1. Marcgrave G, Piso W. Historia Naturalis Brasiliae: In qua non tantum plantæ et animalia, sed et indigenarum morbi, ingenia et mores describuntur et iconibus supra quingentas illustrantur [Internet]. Historia naturalis Brasiliae, : auspicio et beneficio illustriss. I. Mauritii Com. Nassau … adornata : in qua non tantum plantæ et animalia, sed et indigenarum morbi, ingenia et mores describuntur et iconibus supra quingentas illustrantur. Lugdun. Batauorum, Amstelodami: Apud Franciscum Hackium [Leiden], Apud Lud. Elzevirium [Amsterdam]; 1648 [cited 2022 Sep 15]. Available from: http://hdl.handle.net/1887.1/item:1535938

2. Piso W. De India Utriusque Re Naturali et Medica. Amsterdam: Apud Ludovicum et Danielem Elzevirios [Amsterdam]; 1658.

3. Marcgrave G. Historia Rerum Naturalium Brasiliae. In: Historia Naturalis Brasiliae [Internet]. Lugdun. Batauorum, Amstelodami: Apud Franciscum Hackium [Leiden], Apud Lud. Elzevirium [Amsterdam]; 1648 [cited 2022 Sep 15]. p. 50–293. Available from: http://hdl.handle.net/1887.1/item:1535938

4. Whitehead PJP, Boeseman M. A portrait of Dutch 17th century Brazil. Animals, plants and people by the artists of Johan Maurits of Nassau. Amsterdam: North-Holland Publishing Company; 1989.

5. Almeida AV De. Historiae Rervm Naturalivm: ensaios histórico-culturais sobre as ciências biológicas. Recife; 2016.

6. Pickel BJ. Flora do Nordeste do Brasil segundo Piso e Marcgrave no século XVII. Almeida AV De, editor. Recife: EDUFRPE; 2008.

7. BL Sloane MS 1554. Sloane 1554 Chartaceus, in folio, ff.81. sec XVII.Excerpta, ut testatur in Catalogo MS. veteri Johannes Ward, L.L.D., ex Georgii Marggravii seu Maargravii, Brasiliae Historia Naturali, Johannis de Laet, qui opus istud primum edidit manu descripta; cum 17th century.

8. Andrade-Lima D, Maule AF, Pedersen TM, Rahn K. Marcgrave’s Brazilian Herbarium, collected 1638–44. Bot Tidsskr. 1977;71:121–60.

9. Zuidervaart HJ, Matsuura OT. Astronomer, Cartographer and Naturalist of the New World The life and scholarly achievements of Georg Marggrafe (1610-1643) in colonial Dutch Brazil. 2022 Oct.

10. Mentzel C. Prefacio. Theatrum rerum naturalium brasiliae: Brasil-Holandês. In: Ferrão c, Soares JPM, Teixeira DMT, editors. Dutch Brazil. Rio de Janeiro: Editora Index; 1993.

11. Albertin PJ. Arte e ciência no Brasil holandês Theatri Rerum Naturalium Brasiliae: um estudo dos desenhos. Rev Bras Zool. 1985;3:249–326.

12. Whitehead PJP. The original drawings for the Historia naturalis Brasiliae of Piso and Marcgrave (1648). J Soc Bibliogr Nat Hist. 1976;7(4).

13. Lichtenstein MHK. Die Werke von Marcgrave und Piso über die Naturgeschichte Brasiliens, erläutert aus den wieder aufgefundenen Original[zeichnungen]-Abbildungen. In: Isis. 1819.

14. Lichtenstein MHK. Estudo Crítico dos Trabalhos de Marcgrave e Piso sobre a História Natural do Brasil à Luz dos Desenhos Originais. Pinto O, editor. Vol. 2. Brasiliensia Documenta; 1961.

15. Scharf CP. Libri Principis e as ilustrações de fauna do Brasil Holandês: fatura, técnicas, materiais e autores. [Salvador]: Universidade Federal da Bahia; 2019.

16. Brienen RP. Visions of savage paradise: Albert Eckhout, court painter in colonial Dutch Brazil. Amsterdam : Amsterdam University Press; 2006.

17. Teixeira DM. The “Tierbuch” and “Autobiography” of Zacharias Wagener. In: Ferrão c, Soares Jpm, editors. In Dutch Brazil. Rio de Janeiro: Editora Index; 1997.

18. Teixeira DM. A imagem do paraíso: Uma iconografa do Brasil holandês (1624–1654) sobre a fauna e fora do Novo Mundo | The image of paradise: An iconography of Dutch-Brazil (1624–1654) on the New World’s fauna and flora in Brasil-Holandês. Introdução & Miscelânea Cleyeri | Dutch-Brazil. Introduction & Miscellanea Cleyeri. Ferrão c, Soares Jpm, editors. Rio de Janeiro: Editora Index; 1995. 89–183 p.

19. Boeseman M. A hidden early source of information on north-eastern Brazilian zoology.Zool Meded. 1994 Jan 1;68(12):113–25.

20. Whitehead PJP. The biography of Georg Marcgraf (1610-1643/4) by his brother Christian, translated by James Petiver. Soc Bibliogr Nat Hist. 1979;9(3):301–14.

21. Thomsen T. Albert Eckhout: ein niederländischer maler, und sein gönner, Moritz der Brasilianer; ein kulturbild aus dem 17. Jahr-hundert. Levin og Munksgaard; 1938.

22. Joppien R. The Dutch Vision of Brazil. Johan Maurits and his artists. In: Boogaart E, editor. Johan Maurits van Nassau-Siegen 1604-1679 Essays on the occasion of the tercentenary of his death. The Hague: The Johan Maurits van Nassau Stichting; 1979. p. 297–376.

23. Whitehead PJP. Appendix 1: The Marcgrave drawings. In: The clupeoid fishes of the Guianas. Bulletin of the British Museum (Natural History) Zoology; 1973. p. 187–204.

24. Brienen RP. From Brazil to Europe: The zoological drawings of Albert Eckhout and Georg Marcgraf. In: Enenkel KAE, Smith PJ, editors. Early modern zoology: The construction of animals in science, literature and the visual arts. Brill; 2007. p. 273–314.

25. Boeseman M, Holthuis L, Hoogmoed M, Smeenk C. Seventeenth century drawings of Brazilian animals in Leningrad. Zool Verh Leiden. 1990;267(28):1–189.

26. Schneider A. Die Vogelbilder zur Historia Naturalis Brasiliae des Georg Marcgrave. J für Ornithol. 1938 Jan;86(1):74–106.

27. Joost W. Die wundersamen Reisen des Caspar Schmalkalden nach West-und Ostindien 1642-1652. Joost W, editor. Leipzig: Brockhaus Verlag; 1983. 58–64 p.

28. Brienen R. P. Georg Marcgraf (1610–c. 1644): A German cartographer, astronomer, and naturalist-illustrator in Colonial Dutch Brazil. Itinerario. 2001 Mar;25(1):85–122.

29. Gesteira HM. Representations of nature: maps and engravings produced during the Dutch rule in Brazil (1624/1654). Rev do Inst Estud Bras. 2008 Feb;1(46):165–78.

30. Alcantara-Rodriguez M, Françozo M, van Andel T. Plant Knowledge in the Historia Naturalis Brasiliae (1648): Retentions of Seventeenth-Century Plant Use in Brazil. Econ Bot. 2019 Sep;73(3):390–404.

31. Alcantara-Rodriguez M, Françozo M, Van Andel T. Looking into the flora of Dutch Brazil: botanical identifications of seventeenth century plant illustrations in the Libri Picturati. Sci Rep. 2021 Dec 1;11(1).

32. Piso W. De Medicina Brasiliensi. In: Historia Naturalis Brasiliae [Internet]. Lugdun. Batauorum, Amstelodami: Apud Franciscum Hackium [Leiden], Apud Lud. Elzevirium [Amsterdam]; 1648 [cited 2022 Sep 15]. p. 50–120. Available from: http://hdl.handle.net/1887.1/item:1535938

33. Hernández F. Nova plantarum, animalium et mineralium Mexicanorum historia. Blasij Deuersini, Zanobij Masotti, Vitalis Mascardi, editors. Roma: Romae MDCLI; 1651.

34. Monardes NB. De simplicibus medicamentis ex occidentali India delatis quorum in medicina usus est. Antwerp: Plantin; 1574.

35. Clusius C. Rariorum plantarum historia :quae accesserint, proxima pagina docebit. Antwerp: Plantin; 1601.

36. Clusius C. Exoticorvm libri decem :quibus animalium, plantarum, aromatum, aliorum que peregrinorum fructuum historiae describuntur /item Petri Bellonii observationes ; eodem Carolo Clusio interprete ; series totius operis post praefationem indicabitur. Antwerp: Plantin; 1605.

37. Clusius C. Cvrae posteriores, sev Plurimarum non antè cognitarum, aut descriptarum stirpium, peregrinorum’que aliquot animalium novae descriptiones: Quibus & omnia ipsius Opera, aliáque ab eo versa augentur, aut illustrantur [Internet]. Lugduni Batavorum: Plantin; 1611 [cited 2022 Aug 2]. Available from: https://bibdigital.rjb.csic.es/records/item/13494-redirection

38. Ubrizsy A, Heniger J. Carolus Clusius and American Plants. Source: Taxon [Internet]. 1983 Aug [cited 2022 Aug 2];32(3):424–35. Available from: https://www.jstor.org/stable/1221499

39. Piso W. História natural do Brasil [1648]. Correia A, editor. São Paulo: Imprensa Oficial do Estado; 1948.

40. De Laet J. Americae utriusque Descriptio Novus orbis seu Descriptionis Indiae Occidentalis [Internet]. Elzevier; 1633 [cited 2022 Aug 2]. Available from: https://archive.org/details/bub_gb_xLF8OiAonjIC/page/n9/mode/1up

41. De Laet J. L’histoire du nouveau monde ou Description des Indes Occidentales: contenant dix-huict liures [Internet]. Leiden: Elzevier; 1640 [cited 2022 Aug 2]. Available from: https://archive.org/details/lhistoiredunouve00laet/page/n3/mode/2up

42. Barlaeus C. Rerum per octennium in Brasilia et alibi nuper gestarum sub præfectura illustrissimi Comitis I. Mauritii, Nassoviæ, &c. comitis, nunc Vesaliæ gubernatoris & equitatus fderatorum Belgii ordd. sub Auriaco ductoris, historia. [Internet]. Amsterdam: Ex typographeio Ioannis Blaeu; 1647 [cited 2022 Aug 2]. Available from: https://archive.org/details/casparisbarlirer01baer/page/9/mode/thumb

43. Piso W. História Natural, e Médica da índia Ocidental. Leal ML, Carneiro F, Rodrigues E, Rodrigues JH, editors. Rio de Janeiro: Ministério da Educacão e Cultura, Instituto Nacional do Livro; 1957.

44. Piso W. Historia Naturalis & Medicae. Liber quartus: De Arboribus, fructibus, & herbis medicis…. In: De Indiae Utriusque re Naturali et Medica. Amsterdam: Apud Ludovicum et Danielem Elzevirios; 1658. p. 107–266.

45. Piso W. Historia Naturalis & Medicae. Liber quintus: De Noxiis & venenatis, corumque Antiootis. In: De Indiae Utriusque re Naturali et Medica. Amsterdam: Apud Ludovicum et Danielem Elzevirios; 1658. p. 302–19.

46. Brunfels O, Weiditz H. Herbarum vivae eiconeb [sic] ad naturę imitationem, summa cum diligentia et artificio effigiatę, unà cum effectibus earundem, in gratiam veteris illius, & jamjam renascentis herbariae medicinae … Quibus adjecta ad calem, appendix isagogica de usu & administratione simplicium … Argentorati: Apud Joannem Schottum; 1530.

47. Fuchs L. De historia stirpium commentarii insignes, maximis impensis et vigiliis elaborati, adiectis earundem viuis plusquam quingentis imaginibus, numquam antea ad naturae imitationem artificiosius estinctis & expressis [Internet]. Basel: Isingrin; 1542 [cited 2022 Aug 2]. Available from: https://archive.org/details/bub_gb_1uALRWuzDAoC/page/n30/mode/thumb

48. Matthioli PA. New Kreüterbuch : mit den allerschönsten und artlichsten Figuren aller Gewechss, dergleichen vormals in keiner Sprach nie an Tag kommen [Internet]. Venetia: Georgen Melantrich von Auentin, Vincenti Valgriss; 1563 [cited 2022 Aug 2]. Available from: https://www.biodiversitylibrary.org/item/37059#page/14/mode/1up

49. Monardes N. Primera y segvnda y tercera partes dela Historia Medicinal: delas cosas que se traen de nuestras Indias Occidentales, que siruen en Medicina [Internet]. Sevilla: En Sevilla : En casa de Fernando Diaz; 1580. Available from: https://bibdigital.rjb.csic.es/idurl/1/13636

50. Dodoens R. Remberti Dodonaei … trium priorum de stirpium historia commentariorum imagines ad viuum expressae :una cum indicibus, graeca, latina, officinarum, germanica, brabantica, gallicaq[ue] nomina complectentibus. Antverpiae: ex officina Ioannis Loei; 1553.

51. Dodoens R. Stirpium historiae pemptades sex [Internet]. Antwerp: Ex officina Christophori Plantini; 1583 [cited 2022 Sep 14]. Available from: https://archive.org/details/mobot31753000817947/mode/thumb

52. L’Obel M de. Stirpium adversaria nova, perfacilis vestigatio luculentaque accessio ad priscorum, praesertim Dioscoridis, et recentiorum materiam medicam. Quibus propediem accedet altera pars [Internet]. London: T. Purfoot; 1571 [cited 2022 Aug 2]. Available from: https://archive.org/details/b30333180/page/16/mode/2up?form=MY01SV&OCID=MY01SV&form=MY01SV&OCID=MY01SV

53. L’Obel M de. Icones stirpium, seu, Plantarum tam exoticarum, quam indigenarum :in gratiam rei herbariae studiosorum in duas partes digestae : cum septem linguarum indicibus, ad diuersarum nationum vsum. Antwerp: Plantin; 1591.

54. Alpini P. Prosperi Alpini De plantis Aegypti liber : In quo non pauci, qui circa herbarum materiam irrepserunt, errores, deprehenduntur, quorum causa hactenus multa medicamenta ad vsum medicin[ae] admodum expetenda, plerisque medicorum, non sine artis iactura, occu. Venetiis: Apud Franciscum de Franciscis Senensem; 1592.

55. Alpini P. De plantis Aegypti liber […] Editio altera. Patavii: Pauli Frambotti Bibliopolae; 1640.

56. Worm O. Museum Wormianum, seu, Historia rerum rariorum : tam naturalium, quam artificialium, tam domesticarum, quam exoticarum, quae Hafniae Danorum in aedibus authoris servantur [Internet]. Lugduni Batavorum: Iohannem Elsevirium; 1655 [cited 2022 Aug 2]. Available from: https://archive.org/details/gri_museumwormia00worm/page/n185/mode/2up

57. Norton M. Tasting Empire: Chocolate and the European Internalization of Mesoamerican Aesthetics. Am Hist Rev. 2006 Jun 1;111(3):660–91.

58. Donkin RA. Spanish Red: An Ethnogeographical Study of Cochineal and the Opuntia Cactus. Trans Am Philos Soc. 1977 Sep;67(5):1–84.

59. Egmond F. The exotic world of Carolus Clusius: natural history in the making, 1526-1609. In: Ommen K V., editor. The Exotic World of Carolus Clusius (1526-1609) Catalogue of an exhibition on the quatercentenary of Clusius’ death. Leiden: Leidse Universiteitsbibliotheek; 2009.

60. Soares Neto RL, Magalhães FÁL, Tabosa FRS, Moro MF, Costa e Silva MB, Loiola MIB. Flora do Ceará, Brasil: Capparaceae. Rodriguésia. 2014 Sep;65(3):671–84.

61. Margócsy D. Commercial Visions Science, Trade, and Visual Culture in the Dutch Golden Age. University of Chicago Press ; 2014.

62. Nickelsen K. Draughtsmen, botanists and nature: constructing eighteenth-century botanical illustrations. Stud Hist Philos Sci Part C Stud Hist Philos Biol Biomed Sci. 2006 Mar;37(1):1–25.

63. Pardo-Tomás J. Two Glimpses of America from a Distance: Carolus Clusius and Nicolás Monardes. In: Egmond F, Hoftijzer G, Visser RW, editors. Carolus Clusius: Towards a Cultural History of a Renaissance Naturalist. Amsterdam: Koninklijke Nederlandse Akademie van Wetenschappen; 2007.

64. Alcantara-Rodriguez M. Medicinal and other useful plants from Historia Naturalis Brasiliae (1648): Are they currently used in Brazil? [Utrecht]: University of Utrecht– Naturalis Biodiversity Center; 2015.

65. Martius KFP von. Systema materiae medicae vegetabilis Brasiliensis. Frid. Fleischer, Lipsiae; Frid. Beck, Vindobonae; 1843.

66. Schiebinger L., Swan C. Colonial botany: science, commerce, and politics in the early modern world. Schiebinger L., Swan C., editors. University of Pennsylvania Press; 2007.

67. Wijnands DO. The Hortus Medicus Amstelodamensis — its role in shaping taxonomy and horticulture. Kew Mag. 1987 May;4(2):78–91.

68. Linnaeus C. Systema naturae. Stockholm: Holmiae (Laurentii Salvii); 1758.

69. NL-HaNA 1.05.01.01, OWIC, inv.no.58 doc. 206. WIC - National Archives in Den Haag (NL-HaNA, 1.05.01.01, OWIC, inv.no.58 doc. 206.

70. Van den Boogaart E, Brienen RP. Informações do Ceará de Georg Marcgraf (junho-agosto de 1639) in Brasil Holandês. Ferrão C, Soares Jpm, editors. Index; 2002.

71. Monteiro C. Colonial Representations of Brazil and their Current Display in Western Museums: The Mauritshuis Case and the Dutch Gaze. [Leiden]: Leiden University; 2019.

72. Klooster W. The Dutch Moment: War, Trade, and Settlement in the Seventeenth-Century Atlantic World. Cornell University Press; 2016.

73. Fleischer A. Traveling Salesmen or Scholarly Travelers? Early Modern Botanists on the Move Marketing Their Knowledge of Nature. In: Goeing A-S, Parry G, Feingold M, editors. Early Modern Universities. Brill; 2020. p. 371–91.

74. Kusukawa S. The sources of Gessner’s pictures for the Historia animalium. Ann Sci. 2010 Jul 15;67(3):303–28.

75. Kusukawa S. Patron’s review. The role of images in the development of Renaissance natural history. Arch Nat Hist. 2011 Oct;38(2):189–213.

76. Chen JW-H. A Woodblock’s Career. Transferring Visual Botanical Knowledge in the Early Modern Low Countries. Nuncius. 2020 Apr 23;35(1):20–63.

77. Martins L, Fonseca-Kruel VSD, Cabalzar A, Azevedo DL, Milliken W, Nesbitt M, et al. A maloca entre artefatos e plantas: guia da coleção Rio Negro de Richard Spruce em Londres [Internet]. São Paulo: Instituto Socioambiental; 2021 [cited 2022 Aug 2]. Available from: https://acervo.socioambiental.org/acervo/publicacoes-isa/maloca-entre-artefatos-e-plantas-guia-da-colecao-rio-negro-de-richard-spruce

78. Kruel VF, Martins L, Nesbitt M, Milliken W, Coelho-Ferreira M. Nova Pesquisa Sobre as Coleções de Richard Spruce na Amazônia: uma Colaboração Brasil - Reino Unido. Ethnoscientia. 2018 Jul 1;3(2).

79. Cabalzar A, Fonseca-Kruel VSD, Martins L, Milliken W, Nesbitt M. Manual de etnobotânica: plantas, artefatos e conhecimentos indígenas [Internet]. São Paulo: Instituto Socioambiental ; 2017 [cited 2022 Aug 2]. Available from: https://acervo.socioambiental.org/sites/default/files/publications/Manual_de_Etnobotanica_baixa.pdf

